# The HMCES DNA-protein cross-link functions as a constitutive DNA repair intermediate

**DOI:** 10.1101/2021.07.29.454365

**Authors:** Daniel R. Semlow, Victoria A. MacKrell, Johannes C. Walter

## Abstract

The HMCES protein forms a covalent DNA-protein cross-link (DPC) with abasic (AP) sites in ssDNA, and the resulting HMCES-DPC is thought to suppress double-strand break formation in S phase. However, the dynamics of HMCES cross-linking and whether any DNA repair pathways normally include an HMCES-DPC intermediate remain unknown. Here, we show that an HMCES-DPC forms efficiently on the AP site generated during replication-coupled DNA interstrand cross-link (ICL) repair. We use this system to show that HMCES cross-links form on DNA after the replicative CMG helicase has passed over the AP site, and that HMCES is subsequently removed by the SPRTN protease. The HMCES-DPC suppresses DSB formation, slows translesion synthesis (TLS) past the AP site, and introduces a bias for insertion of deoxyguanosine opposite the AP site. These data show that HMCES-DPCs can form as constitutive intermediates in replication-coupled repair, and they suggest a general model of how HMCES protects AP sites during DNA replication.

## Main

DNA abasic (AP) sites are common genomic lesions that arise from spontaneous DNA depurination/depyrimidination or as intermediates in the excision of damaged nucleobases by DNA glycosylases. When formed in double-stranded DNA, AP sites are usually incised by base excision repair factors (AP endonucleases and AP lyases), generating a one nucleotide gap that is filled in by DNA polymerase *β*^1^. AP sites are also susceptible to spontaneous cleavage through a *β*-elimination mechanism^2^. Importantly, enzymatic or spontaneous cleavage of AP sites located in ssDNA, e.g. during DNA replication, results in the formation of a DNA double-stranded break (DSB), which introduces the potential for gross chromosomal rearrangements.

Recently, Cortez and colleagues demonstrated that the 5-hydroxymethylcytosine binding, embryonic stem cell-specific (HMCES) protein associates with replication forks, covalently cross-links to ssDNA AP sites, and suppresses accumulation of DSBs in cells treated with genotoxic agents that induce AP site formation^3^. Structural evidence indicates that the universally conserved SRAP domain of HMCES specifically cross-links to AP sites positioned in ssDNA *in vitro* through formation of a thiazolidine linkage^4–7^. Taken together, these observations indicate that HMCES stabilizes ssDNA AP sites during replication through formation of a DNA-protein cross-link (HMCES-DPC).

HMCES-DPC formation also appears to suppress mutagenic bypass of AP sites by translesion synthesis (TLS). Loss of HMCES increases association of the REV1-Pol*ζ* TLS polymerase with replication forks and increases the mutagenicity of UV-induced DNA damage^3, 8^. Additionally, HMCES depletion slows replication forks upon induction of AP sites, and this defect is reversed by inhibition of REV1-Pol*ζ* or depletion of the TLS polymerase Pol*κ*^9^. These data suggest that HMCES antagonizes AP site bypass by TLS polymerases and promotes a more rapid, error-free bypass mechanism, possibly involving template switching.

Despite these advances, it is unclear how formation of the HMCES-DPC is coupled to DNA replication. HMCES-DPC formation has so far been examined only in the context of (1) cellular treatment with genotoxic agents that indirectly induce AP site formation by introducing a spectrum of DNA damage^3, 8^ or (2) ectopic expression of APOBEC3A, which deaminates deoxycytosine to deoxyuracil that is in turn converted to an AP sites by uracil DNA glycosylase^9, 10^. In the former case, the identity of the DNA lesions that are encountered during replication and trigger HMCES action remain unknown. In the latter case, HMCES accumulates on chromatin independently of replication^9^, leaving open the mechanism by which these HMCES-DPCs form. In both cases, the orientation of the relevant DNA lesions relative to fork progression cannot be distinguished; therefore, it is unclear whether HMCES-DPCs form mainly on the lagging strand template, as previously suggested^3, 9^, or also on the leading strand template. Another important question is whether HMCES-DPCs are only formed *ad hoc* when a replication fork encounters a spontaneously-formed AP site, or whether they are generated constitutively as part of any DNA repair pathways that involve an AP site intermediate, in which case an efficient mechanism would be required to remove the HMCES-DPC after it has served its function.

AP sites can also react with exocyclic amines of nucleobases on the opposite strand to generate highly toxic DNA interstrand cross-links (ICLs)^11, 12^. These AP site-induced ICLs (AP-ICLs) covalently link the two strands of DNA and block DNA replication and transcription. During replication, AP-ICLs are repaired by the NEIL3 DNA glycosylase^13^. This pathway is activated when the convergence of two replication forks on either side of the ICL stimulates ubiquitylation of the replicative CDC45-MCM2-7-GINS (CMG) helicase by the E3 ubiquitin ligase TRAIP (Fig. 1a and ref.^14^). NEIL3 is recruited to ubiquitylated CMGs and unhooks the ICL by cleaving an *N*-glycosyl bond in the lesion, regenerating an AP site. The leading strand of one fork encounters the AP site and is then extended past the lesion in a REV1-dependent manner^13^. If the NEIL3 pathway fails to unhook the ICL, the ubiquitin chains on CMG are extended and the helicase is unloaded by the p97 segregase^15^. At this point, the Fanconi anemia (FA) pathway unhooks the ICL through nucleolytic incisions, converting the ICL into a DSB that is subsequently repaired by homologous recombination^16–18^. Significantly, the NEIL3 pathway is prioritized over the FA pathway, likely because the NEIL3 pathway avoids formation of a DSB intermediate and thus minimizes the potential for genome instability^14, 19^. NEIL3 does, however, introduce a labile ssDNA AP site, the fate of which is unknown.

**Fig. 1:**
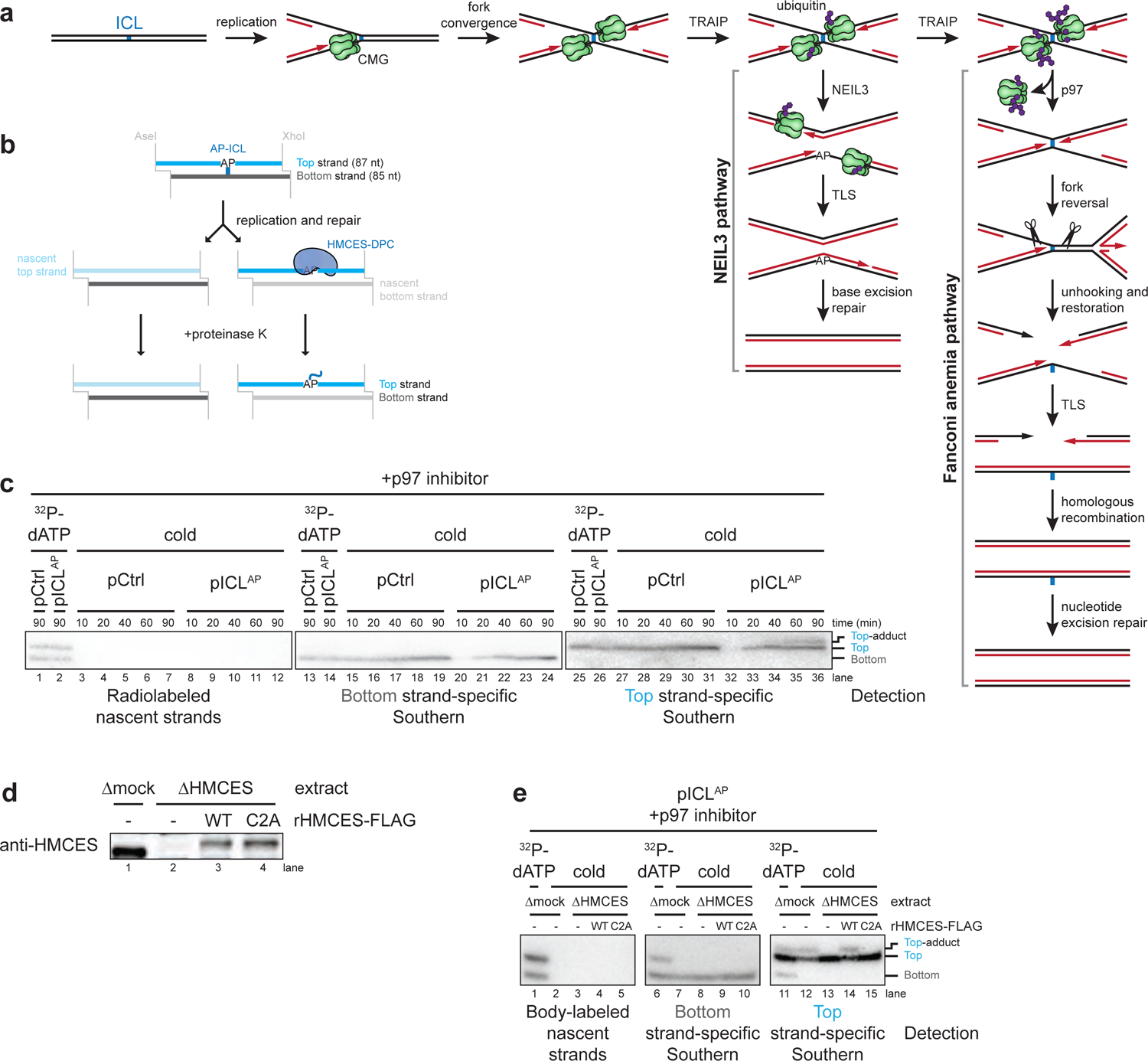
HMCES cross-links to AP sites during NEIL3-dependent ICL repair. **a**, Model of replication-coupled ICL repair pathways. **b**, Schematic of repair products produced by AseI and XhoI digestion. AseI and XhoI digestion allows resolution of the top (87 nt) and bottom (85 nt) strands. AP-ICL unhooking by NEIL3 generates an AP-site in the top parental strand that cross-links to HMCES and generates a discrete adduct after proteinase K digestion. **c**, Detection of AP-site adducts by strand-specific Southern blotting. pCtrl or pICL^AP^ was replicated in egg extract. The p97 inhibitor NMS-873 was added to prevent activation of the FA pathway. At the indicated times, DNA was isolated, treated with proteinase K, and digested with AseI and XhoI. The repair products were separated on a denaturing polyacrylamide sequencing gel and visualized by Southern blotting with the indicated strand-specific probes. Size markers were generated by replicating pCtrl and pICL^AP^ in extracts supplemented with [*α*-^32^P]dATP, which generates radiolabeled nascent strands that were similarly processed and resolved on the same sequencing gel. **d**, HMCES immunodepletion. The extracts used in the replication reactions shown in **e** were blotted for HMCES. **e**, pICL^AP^ was replicated using the extracts in **d** and supplemented with p97 inhibitor and rHMCES, as indicated. Repair products were visualized by body labeling or strand-specific Southern blotting as in **c**.

Here, we used *Xenopus* egg extracts to show that an HMCES-DPC efficiently forms during AP-ICL repair. We exploited this system to study the dynamics of HMCES cross-linking to DNA and its subsequent removal. We find that after replisome stalling at the AP-ICL and ICL unhooking by NEIL3, CMG translocates beyond the newly generated AP-site followed by HMCES reaction with the AP site. The leading strand then collides with the HMCES-DPC, which triggers DPC degradation by the replication-coupled protease SPRTN. Functionally, the HMCES-DPC suppresses DSB formation, slows TLS past the AP site, and promotes insertion of deoxyguanosine opposite the AP site. Our results show that HMCES-DPCs can participate as a constitutive intermediate in DNA repair, and they suggest a general model for the generation and resolution of HMCES-DPCs at the DNA replication fork.

## Results

### HMCES forms a DPC during NEIL3-dependent ICL repair

It is presently unknown whether an HMCES-DPC represents a constitutive intermediate in any DNA repair pathway. We speculated that HMCES might cross-link to the AP site generated during NEIL3-dependent ICL repair. To test this idea, we replicated an undamaged control plasmid (pCtrl) or an AP-ICL containing plasmid (pICL^AP^) in *Xenopus* egg extracts. A DNA fragment encompassing the ICL was excised, and the top and bottom strands were resolved on a denaturing polyacrylamide gel and visualized by strand-specific Southern blotting (Fig. 1b).

Prior to electrophoresis, the DNA was treated with proteinase K to convert any DPC formed to a short peptide adduct that is predicted to cause a discrete mobility shift^20^. As expected, replication of both pCtrl and pICL^AP^ yielded only unmodified bottom strands (Fig. 1b and 1c, lanes 15-24). In contrast, while pCtrl replication produced only an unmodified top strand, pICL^AP^ replication produced an additional, slower-migrating top strand species, consistent with the presence of a peptide adduct (Fig. 1c, lanes 27-36). After HMCES depletion (Fig. 1d), the slower-migrating top strand disappeared, and it reappeared with addition of active, recombinant HMCES (rHMCES)(Fig. 1e, lanes 12-14 and Extended Data Fig. 1a,b). In contrast, addition of HMCES protein harboring a C2A mutation (rHMCES^C2A^) that blocks formation of the AP site thiazolidine linkage did not support formation of the slow-migrating top strand (Fig. 1e, lane 15, Extended Data Fig. 1b, and ref.^4^). We conclude that HMCES cross-links to the AP site-containing top strand during AP-ICL repair, leading to the formation of a covalent HMCES-DNA intermediate.

We wanted to determine the efficiency of HMCES-DPC formation. During replication-coupled ICL repair, 50% of the top strands derive from newly synthesized DNA (templated by the parental bottom strand) and 50% derive from the parental, AP site-containing top strand (Fig. 1b). We therefore expect at most 50% of total top strands to be adducted by HMCES. Quantification of Fig. 1c shows that 25.3% of total top strands contained an adduct. Thus, at least 50.6% of parental top strands were cross-linked by HMCES. The actual efficiency of cross-linking might be higher since proteinase K treatment may not convert all HMCES DPCs to a homogenous peptide adduct.

As an orthologous approach to determine the efficiency of HMCES cross-linking, we measured the sensitivity of replication intermediates to digestion with recombinant (r)APE1, which cleaves the phosphodiester backbone 5’ of an AP site, but not when the AP site is cross-linked by HMCES^3^. pICL^AP^ was replicated in mock- or HMCES-depleted extract and repair intermediates were recovered and digested with HincII and rAPE1 (Extended Data Fig 1c).

Since NEIL3-dependent unhooking introduces an AP site in only one of the two daughter plasmids, we expected no more than 50% of plasmids to be cleaved by rAPE1 in the absence of HMCES^13^. At the 45-minute time point, only a fraction of plasmids recovered from mock-depleted extract was cleaved upon treatment with rAPE1 (13.9% of total plasmids; 27.8% of AP site-containing plasmids; Extended Data Fig. 1d-f). Strikingly, HMCES-depletion resulted in a dramatic increase in the fraction of plasmids cleaved by rAPE1 (49.6% of total plasmids; 99.2% of AP site-containing plasmids; Extended Data Fig. 1d-f). This cleavage was suppressed by the addition of rHMCES, but not rHMCES^C2A^. Inferring that the difference in rAPE1 cleavage efficiency between samples recovered from mock- and HMCES-depleted extract reflects the efficiency of HMCES-DPC formation, we estimate that HMCES-DPCs formed on ∼71.4% (99.2% − 27.8%) of AP sites generated during NEIL3-dependent ICL repair. The time-dependent decline in rAPE1 susceptibility likely reflects placement of the AP site into dsDNA by TLS^13^. In conclusion, our results indicate that an HMCES-DPC represents a major intermediate in AP-ICL repair. Furthermore, these data indicate that the endogenous HMCES adduct can protect the AP site from cleavage by exogenously added APE1, consistent with replication-coupled formation of an irreversible thiazolidine linkage.

### HMCES suppresses DSB formation during NEIL3-dependent ICL repair

We then asked how the HMCES-DPC affects AP-ICL repair in egg extracts. To this end, we replicated pICL^AP^ in mock- or HMCES-depleted egg extracts supplemented with [*α*-^32^P]dATP and resolved replication and repair intermediates on a native agarose gel. As shown previously^13^, replication in mock-depleted extract resulted in convergence of replication forks at the ICL, resulting in a slow-migrating “figure 8” intermediate (Fig. 2a, lanes 1-8). Unhooking by NEIL3 then lead to the appearance of catenated daughter molecules, followed by the formation of open circular products that were finally converted to supercoiled species after TLS. HMCES-depletion did not significantly alter the accumulation of replication intermediates or open-circular and supercoiled products (Fig. 2a, lanes 9-16). However, it did cause transient and low-level accumulation of a new, linear species that is consistent with cleavage of the AP site generated during unhooking (Fig. 2a, red arrowhead). Interestingly, disappearance of the linear species correlated with the appearance of radioactive products in the wells (Fig. 2a, blue arrowhead). We suspect that these well products comprise joint molecules that arise as a consequence of homology directed repair of the DSB^17^. Formation of linear species and well products was suppressed by rHMCES but not rHMCES^C2A^ (Fig. 2a, lanes 17-32). Addition of rHMCES^ΔPIP^ harboring amino acid substitutions that disrupt its conserved PCNA-interacting protein-box (PIP-box) motif^3^ also partially suppressed linear species and well product formation (Extended Data Figs. 1a,b and 2b,c), indicating that the HMCES PIP-box is not absolutely required to suppress DSBs during AP-ICL repair in egg extract. Significantly, HMCES-depletion had no effect on replication of pCtrl or a plasmid with a cisplatin-ICL (pICL^Pt^) that is repaired exclusively by the Fanconi anemia repair pathway without formation of an AP site (Extended Data Fig. 2d,e). These results show that HMCES suppresses cleavage of the AP site by endogenous nuclease(s), preventing DSB formation during AP-ICL repair.

**Fig. 2:**
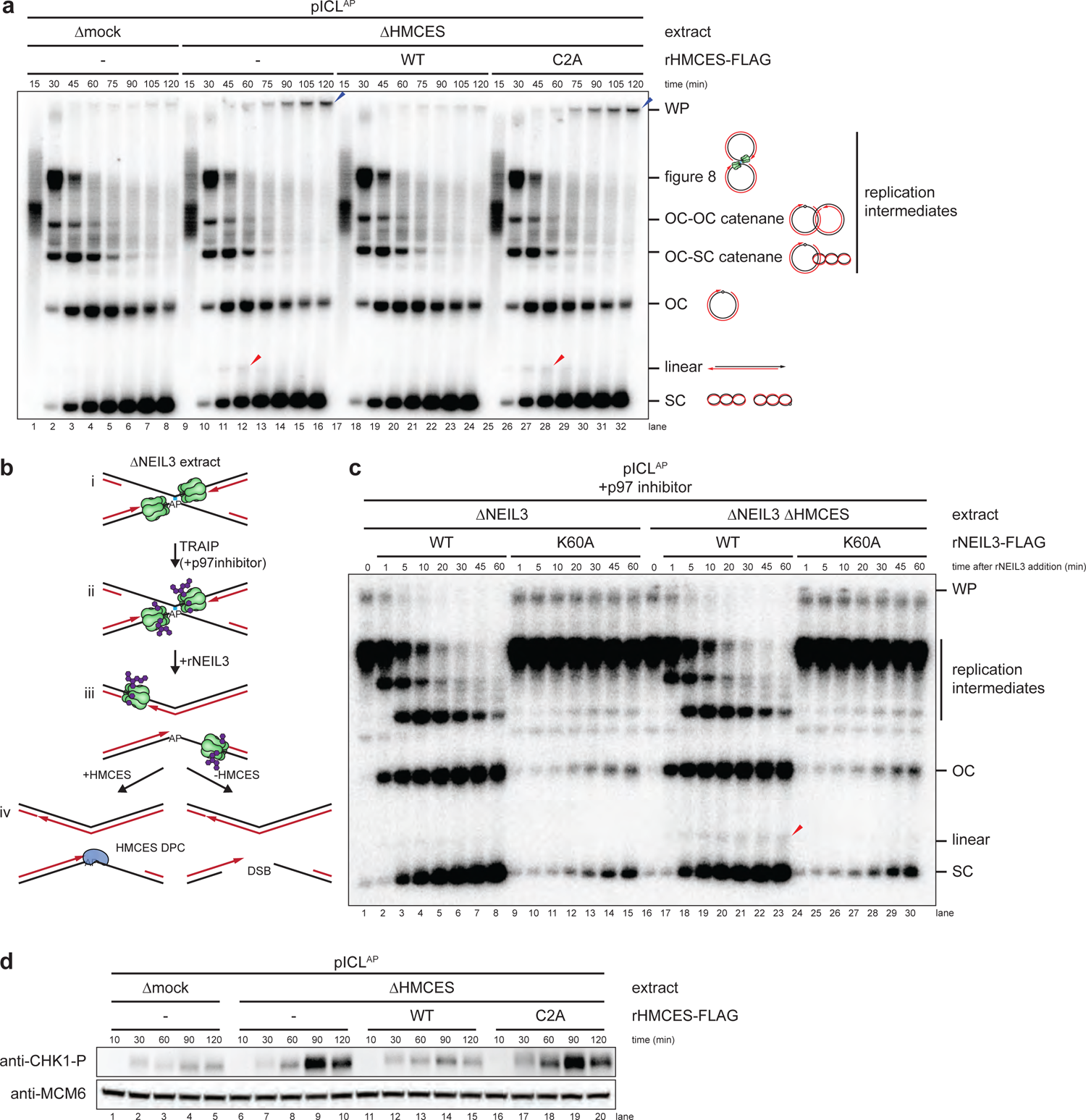
HMCES suppresses DSB formation during NEIL3-dependent ICL repair. **a**, pICL^AP^ was replicated with [*α*-^32^P]dATP in mock- or HMCES-depleted extracts supplemented with rHMCES, as indicated. Replication intermediates were separated on a native agarose gel and visualized by autoradiography. SC, supercoiled; OC, open circular; WP, well products. Red arrowheads indicate linear species; blue arrowheads indicate well-products. **b**, Experimental strategy to synchronize AP-ICL unhooking by NEIL3-depletion and add back. p97 inhibitor is added to replication reactions to prevent CMG unloading and accumulate replication forks that have converged at the ICL. rNEIL3 is added to stalled forks to activate unhooking. **c**, pICL^AP^ was replicated with [*α*-^32^P]dATP as described in (b) using the NEIL3- or NEIL3- and HMCES-depleted extracts shown in Extended Data Fig. 2a. Replication intermediates were analyzed as in **a**. **d**, pICL^AP^ was replicated in mock- or HMCES-depleted egg extracts (shown in Extended Data Fig. 2g) supplemented with rHMCES, as indicated. Replication reactions were separated by SDS-PAGE and blotted for phospho-CHK1 and MCM6 (loading control).

We next addressed whether formation of the linear species observed in HMCES-depleted extract depends on NEIL3 glycosylase activity. To this end, we replicated pICL^AP^ in NEIL3- and HMCES-depleted extract supplemented with the p97 inhibitor NMS-873 which blocks CMG unloading and activation of the FA pathway^13–15^. This experimental setup led to the convergence of replication forks at the ICL but no unhooking (Fig. 2b, i-ii and Extended Data Fig. 2f). To induce unhooking and AP site formation, we then added back wild-type recombinant NEIL3 (rNEIL3) or catalytically defective NEIL3 (rNEIL3^K60A^; ref.^13^) and resolved the DNA on a native agarose gel (Fig. 2b, iii-iv, and Fig. 2c). As expected, addition of rNEIL3, but not rNEIL3^K60A^, resulted in rapid AP-ICL unhooking as indicated by conversion of cross-linked replication intermediates into open-circular and supercoiled products (Fig. 2c, lanes 16-30).

Importantly, addition of rNEIL3, but not rNEIL3^K60A^, generated the linear species indicative of AP site cleavage (Fig. 2c, lanes 16-30). As seen in Fig. 2a, fewer linear species were generated in the presence of HMCES (Fig. 2c, lanes 1-15). We conclude that HMCES suppresses DSB formation during ICL repair by cross-linking to the AP site produced by NEIL3-dependent *N*-glycosyl bond cleavage.

To seek independent evidence for the generation of DNA damage in the absence of HMCES, we replicated pICL^AP^ in mock- or HMCES-depleted egg extracts and measured CHK1 phosphorylation. CHK1 is phosphorylated by ATR, whose activation depends on the formation of single stranded DNA^21^. Replication of pICL^AP^ in HMCES-depleted extract led to a marked increase in ATR-dependent CHK1 phosphorylation compared to mock-depleted extract (Fig. 2d, lanes1-10 and Extended Data Fig. 2h,i), suggesting that after AP site cleavage and DSB formation, DNA ends are resected to generate ssDNA. Addition of wild-type rHMCES, but not rHMCES^C2A^, suppressed CHK1 phosphorylation (Fig. 2d, lanes 11-20). HMCES-depletion did not stimulate CHK1 phosphorylation during replication of pCtrl or pICL^Pt^ (Extended Data Fig. 2j,k). Together, these results provide further evidence that HMCES prevents DNA damage by shielding the AP site generated during AP-ICL repair.

We asked why only a small fraction of AP sites are converted to DSBs in the absence of HMCES (Fig. 2a). One explanation is that the leading strand is usually extended beyond the AP site by TLS before AP site cleavage, placing the AP site in dsDNA and preventing DSB formation. However, this was not the case because REV1 depletion (Extended Data Fig. 3a) did not enhance DSB formation in the absence of HMCES (Extended Data Fig. 3b, lanes 15-28).

**Fig. 3:**
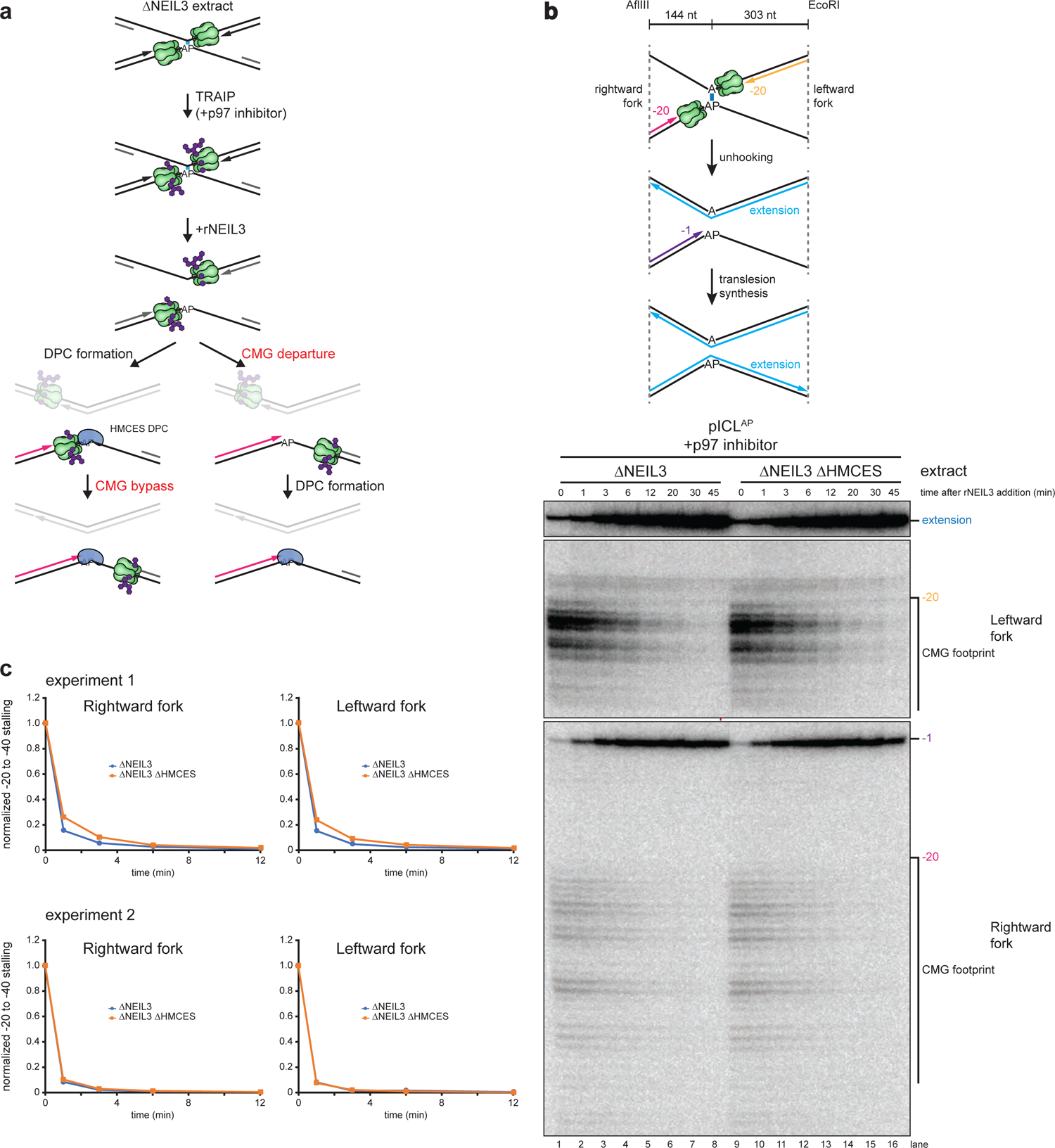
HMCES-DPC formation does not impede CMG translocation past AP site. **a**, Models for timing of HMCES-DPC formation. Left branch, HMCES-DPC formation precedes CMG bypass of AP site, which is predicted to delay rightward leading strand approach to the AP site. Right branch, CMG bypass of AP site precedes HMCES-DPC formation, resulting in no delay in rightward leading strand approach. **b**, Top, Schematic of nascent strands generated during ICL repair. AflIII and EcoRI cut 144 nucleotides to the left and 303 nucleotides to the right of the ICL, respectively, generating characteristic −20 stall, −1 stall, and strand extension products. Bottom, pICL^AP^ was replicated with [*α*-^32^P]dATP and p97 inhibitor in the indicated extracts (shown in Extended Data Fig. 4a) and nascent DNA strands were isolated, digested with AflIII and EcoRI, and resolved by denaturing polyacrylamide gel electrophoresis. Top, middle, and bottom panels show sections of the same gel to visualize extension, leftward leading strands, and rightward leading strands, respectively. **c**, The persistence of −20 to −40 stall products in **b** was quantified by dividing the summed intensities of the −20 to −40 stall product bands in each lane by the intensity of the rightward fork −1 stall product band. Quantifications were normalized to the accumulation of −20 to −40 stall products at the 0 min time timepoint. Quantifications from two independent experiments are shown.

REV1-depletion was successful because it blocked the conversion of open circular plasmid into closed supercoiled plasmid, consistent with a TLS defect (Extended Data Fig. 3b). These results imply the existence of an alternative, redundant mechanism that suppresses AP site cleavage during ICL repair. For example, it is possible that the single-stranded DNA binding protein RPA inhibits AP site cleavage, but this hypothesis is difficult to test because RPA is required for DNA replication^22^. In conclusion, the HMCES-DPC is critical to suppress DSB formation during NEIL3-dependent ICL repair, but in its absence, overlapping mechanisms likely perform the same function.

### Evidence that HMCES cross-links to the AP site after CMG progression

We next addressed when HMCES cross-links to the AP site. After CMGs converge on an ICL and NEIL3 unhooks the lesion, CMGs translocate past the newly generated AP site, and the leading strand advances to and stalls at the −1 position, before undergoing TLS past the lesion (Fig. 1a). One scenario is that HMCES cross-links to the AP site *before* CMG moves beyond the AP site (Fig. 3a, left branch), in which case CMG would likely need to bypass the intact HMCES-DPC, as seen previously for the repair of HpaII-DPCs^23^. Alternatively, HMCES might cross-link to the AP site *after* CMG departure (Fig. 3a, right branch). In the former scenario, HMCES should delay the approach of the nascent leading strand to the −1 position because CMG bypass of an intact DPC is a slow event^23^ (Fig. 3a, left branch). In the latter scenario, HMCES should not affect leading strand approach to −1 (Fig. 3a, right branch). To distinguish between these models, we replicated pICL^AP^ in NEIL3-depleted or NEIL3- and HMCES-depleted extract supplemented with p97 inhibitor, added back rNEIL3 to activate unhooking (Extended Data Fig. 4b), and monitored the kinetics of rightward leading strand extension to the −1 position. Importantly, HMCES depletion had no discernible effect on the timing of leading strand approach to −1 (Fig. 3b,c), showing that CMG translocation past the AP site is unaffected by HMCES. Moreover, the kinetics of nascent strand extension were indistinguishable for the rightward replication forks, which encounter a HMCES-DPC on the template strand, and the leftward forks, which do not (Fig. 3b, compare middle and bottom panels, and Fig. 3c), indicating that CMG translocation is unaffected by the HMCES-DPC. These observations strongly suggest that HMCES cross-links to DNA after CMG has translocated past the AP site.

**Fig. 4:**
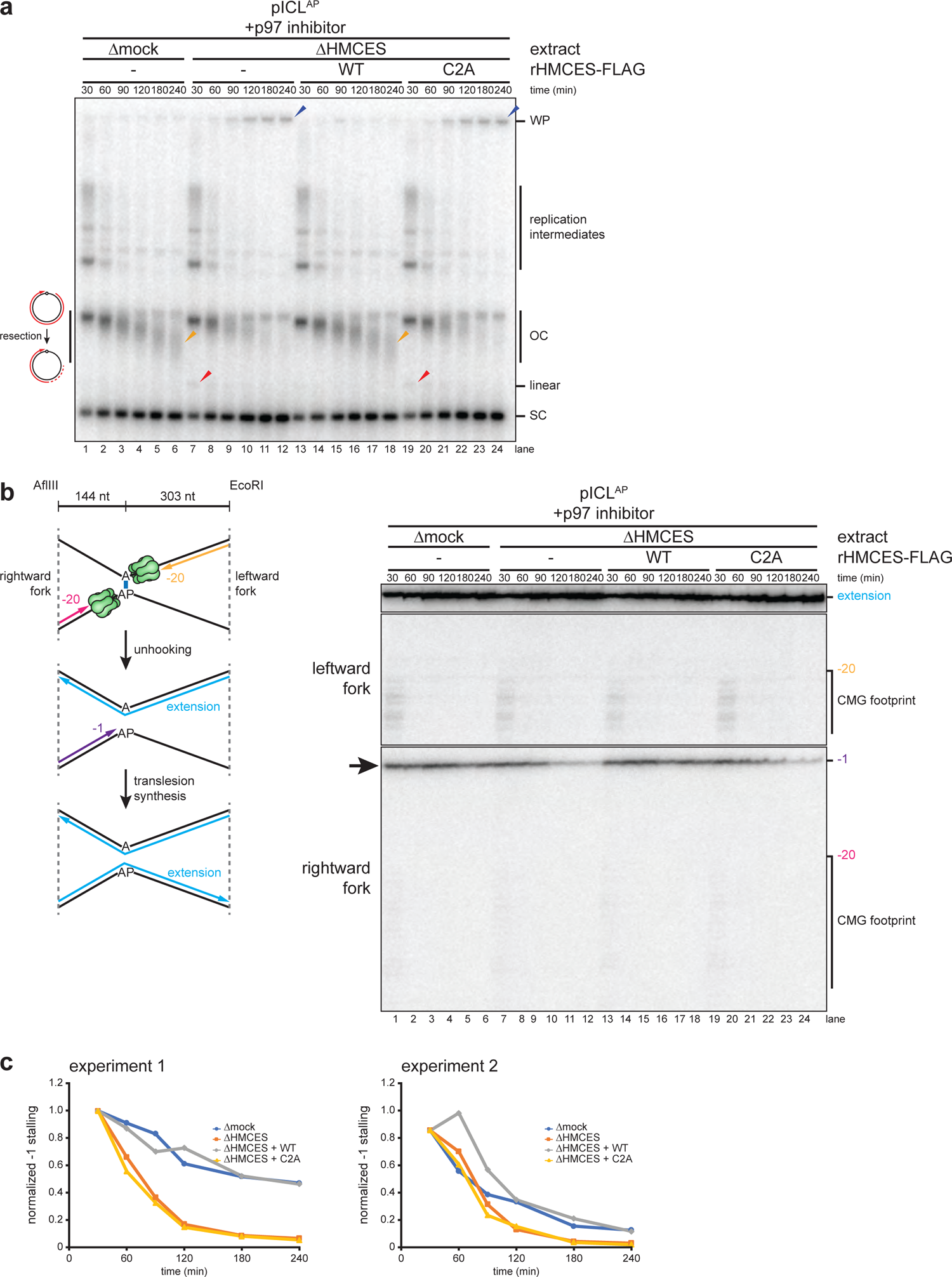
HMCES-DPC formation impedes translesion synthesis. **a**, pICL^AP^ was replicated with [*α*-^32^P]dATP and p97 inhibitor in mock- or HMCES-depleted extracts (shown in Extended Data Fig. 4c) supplemented with rHMCES, as indicated. Replication intermediates were analyzed as in Fig. 2a. Orange arrowheads indicate degraded open circular plasmids. **b**, Nascent DNA strands from the pICL^AP^ replication reactions shown in **a** were analyzed as in Fig. 3b. **c**, The persistence of the rightward fork −1 stall product in **b** was quantified by dividing the intensity of the −1 stall product band in each lane by the intensity of the full length extension product band. Quantifications were normalized to the accumulation of the −1 stall product at the 0 min time timepoint. Quantifications from two independent experiments are shown.

### HMCES-DPC formation impedes TLS past the AP site

We wanted to determine if HMCES impedes TLS past an AP site. To this end, we replicated pICL^AP^ in mock-depleted or HMCES-depleted extract and analyzed nascent strand extension. To suppress the FA pathway, extracts were treated with p97 inhibitor, which also partially inhibits TLS. Under these conditions, replication in mock-depleted extract resulted in a diffuse smear of DNA just below the position of open-circular plasmid, likely due to resection of unligated nascent 5’ ends on plasmids where TLS had not yet occurred (Fig. 4a, lanes 1-6). Interestingly, HMCES-depletion decreased the abundance of the faster migrating open-circular species and increased the amount of supercoiled DNA (Fig. 4a, lanes 7-12), suggesting that TLS past the AP site proceeds more efficiently in the absence of an HMCES-DPC. Consistent with this interpretation, the −1 stall product of the rightward leading strand disappeared more rapidly in HMCES-depleted extract than in mock-depleted egg extract (Fig. 4b, lanes 1-12, black arrow, and Fig. 4c), indicating that the HMCES-DPC impedes TLS past the AP site. Addition of rHMCES, but not rHMCES^C2A^, restored both the accumulation of open-circular species and the delay in nascent strand extension (Fig. 4a-c). Combined with our other observations (Fig. 3), these data indicate that the HMCES-DPC forms after CMG translocates beyond the AP site but before TLS.

### HMCES-DPC formation alters the mutation signature of NEIL3-dependent ICL repair

Given that the HMCES-DPC delays bypass of the AP site during ICL repair, we wondered whether it influences the mechanism and mutagenicity of AP site bypass. We therefore performed next generation sequencing of AP-ICL repair products. AP-ICL plasmids were replicated in mock-depleted extract, HMCES-depleted extract, or HMCES-depleted extract supplemented with rHMCES or rHMCES^C2A^, and next generation sequencing was performed to determine which nucleotide is incorporated across from the AP site in each condition (Fig. 5a and Extended Data Fig. 5a-f). Since bypass of the AP site through a template-switching mechanism should yield products whose sequence is dictated by the nucleotide *opposite* the AP site^24^ (Extended Data Fig. 5a), we also examined the effect of varying the base opposite the AP site. For each extract condition, we obtained >11,000 reads that were derived from the nascent DNA strand produced upon bypass of the AP site (Extended Data Fig. 5b, orange box).

**Fig. 5:**
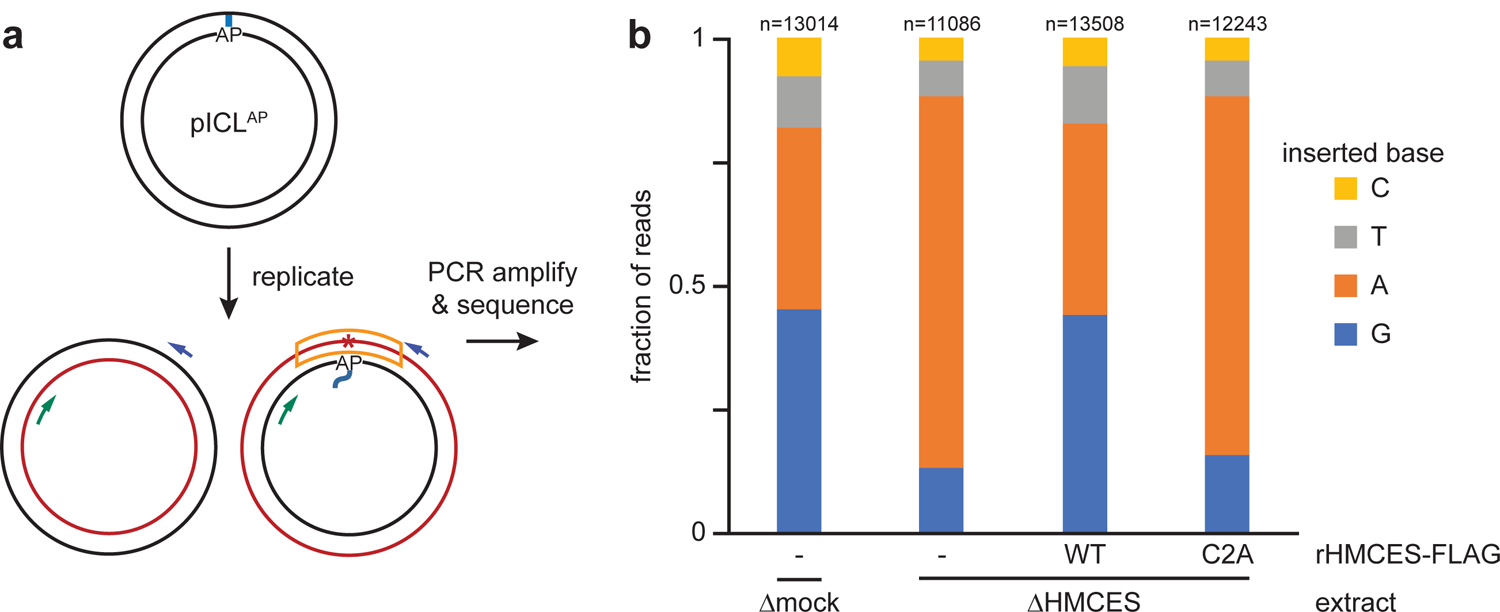
The HMCES-DPC promotes dG insertion during translesion synthesis. **a**, Diagram depicting the strand that is sequenced after NEIL3-dependent ICL repair. Blue and green arrows indicate PCR primers; asterisk indicates potential point mutation; the sequenced strand is boxed in orange. See Extended Data Fig. 5b for details of experimental strategy and plasmid design. **b**, The four pICL^AP^ plasmids described in Extended Data Fig. 5b were pooled and replicated using the indicated extracts (shown in Extended Data Fig. 5d), and sequencing libraries were prepared as described in Extended Data Fig. 5b. The fraction of reads corresponding to insertion of a given nucleotide opposite the AP site is plotted for each extract condition. n, number of pooled nascent strand reads obtained for each condition (see Extended Data Fig. 5g for deconvolution of pooled reads). Next generation sequencing read counts can be found in Supplementary Table 2.

Importantly, in all conditions, the identity of the nucleobase opposite the AP-ICL did not influence which base was inserted (Extended Data Fig. 5g), indicating that TLS, and not template switching, is the dominant mode of AP site bypass in egg extracts. In mock-depleted extract, the purines dG and dA were the most common nucleotides inserted opposite the AP site (45.3% for dG, 36.6% for dA), while the pyrimidines dT (10.5%) and dC (7.7%) were inserted much less frequently (Fig. 5b). By contrast, in HMCES-depleted extract, dA comprised 75.2% of reads while dG dropped to 13.0% (Fig. 5b). Pyrimidines were still inserted in only a minor fraction of bypass events in the absence of HMCES (7.2% for dT and 4.6% for dC).

Addition of wild-type rHMCES, but not rHMCES^C2A^, to HMCES-depleted extract restored the proportion of dG insertions to levels observed in mock-depleted extract (Fig. 5b). These results indicate that while the REV1-dependent TLS prefers to insert dA opposite an unprotected AP site (consistent with the “A rule” for TLS polymerases), formation of an HMCES-DPC biases the insertion step in favor of dG incorporation. Overall, our results implicate dG point mutation as a characteristic signature of HMCES-DPC bypass by TLS.

### The SPRTN protease promotes HMCES-DPC removal

Finally, we asked how HMCES is removed from DNA following cross-linking to the AP site during ICL repair. Previous work indicated that either the proteasome or the SPRTN protease can promote replication-coupled degradation of a HpaII DPC^20, 25^. We therefore tested how these proteases contribute to removal of the HMCES-DPC. To this end, we replicated pICL^AP^ in NEIL3-depleted or NEIL3- and SPRTN-depleted extract, which stalls replication forks on either side of the ICL (Fig. 6a,b). We then added back rNEIL3 to activate unhooking and recovered chromatin to monitor HMCES association by immunoblotting. Compared to the mock-depleted control, SPRTN depletion led to robust accumulation of HMCES on chromatin (Fig. 6c, top panel, compare lanes 3-5 and 11-13). In contrast, treatment with the proteasome inhibitor MG262 had no effect, either alone or in combination with SPRTN depletion (Fig. 6c, top panel). Efficient removal of HMCES from chromatin was restored in SPRTN-depleted extract by addition of wild-type recombinant SPRTN (rSPRTN^WT^), but not catalytically inactive SPRTN (rSPRTN^E89Q^; ref.^25^; Extended Data Fig. 6a,b). These results indicate that SPRTN is the major protease that degrades HMCES-DPCs. Importantly, HMCES association with chromatin was not detected during replication of pCtrl or pICL^Pt^ (Extended Data Fig. 6c), providing further evidence that stable association of HMCES with DNA requires an AP site. Interestingly, in the absence of SPRTN, HMCES accumulated as a diffuse smear of higher molecular weight species that collapsed into a single HMCES band upon treatment with the general deubiquitylating enzyme rUSP21 (Fig. 6c, compare top and bottom panels). This observation indicates that HMCES is extensively polyubiquitylated, although the failure of MG262 to stabilize HMCES in the presence or absence of SPRTN suggests that this ubiquitylation does not target HMCES to the proteasome.

**Fig. 6:**
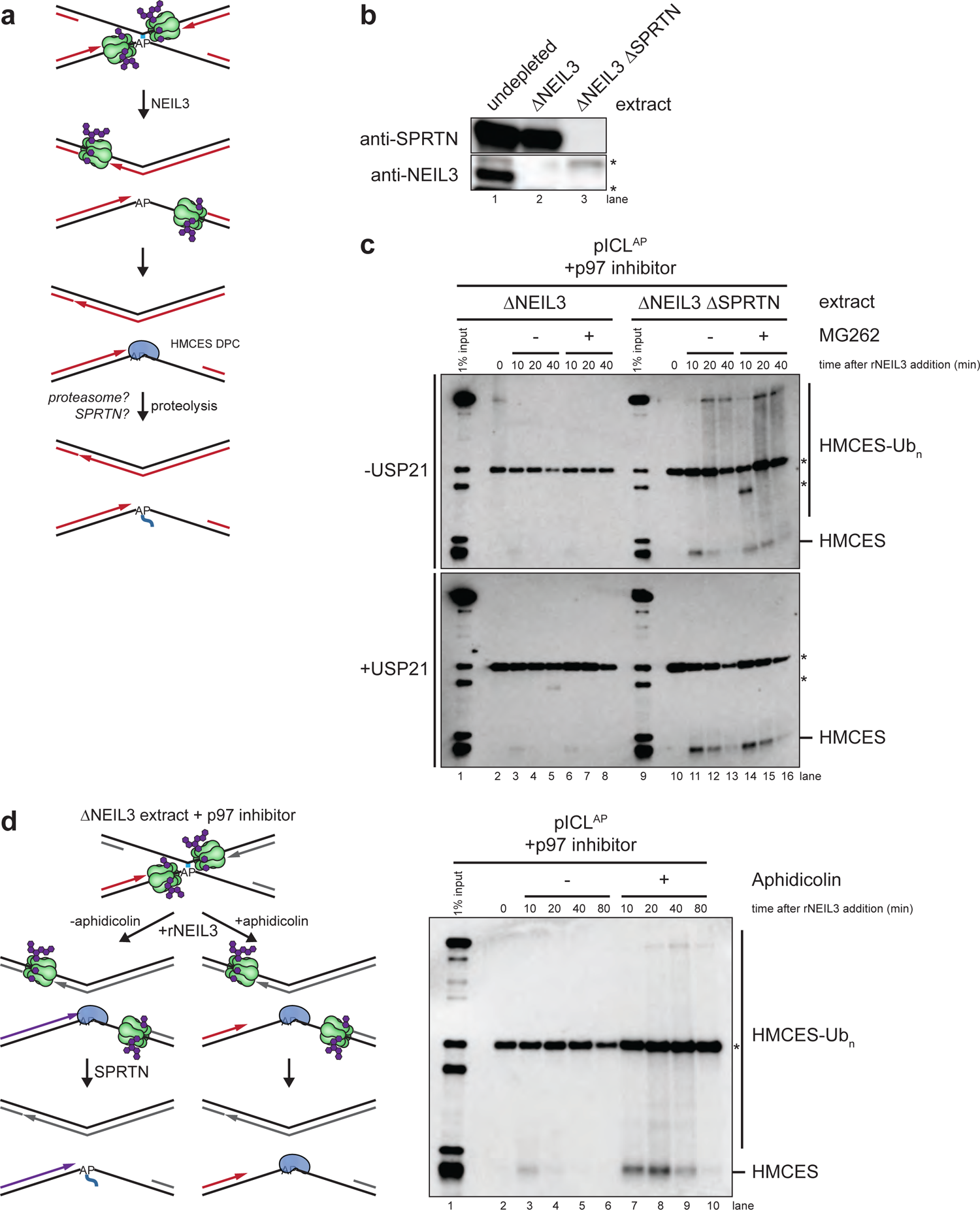
SPRTN degrades HMCES-DPCs formed during ICL repair. **a**, Schematic of HMCES-DPC proteolysis during ICL repair. **b**, NEIL3 and SPRTN immunodepletion. The extracts used in the replication reactions shown in **c** were blotted for NEIL3 and SPRTN. Non-specific bands are marked with asterisks. **c**, pICL^AP^ was replicated in the indicated egg extracts supplemented with p97 inhibitor for 60 min. Reactions were then treated with the proteasome inhibitor MG262 (or DMSO control) and rNEIL3 to promote ICL unhooking. Chromatin was recovered under stringent conditions, and associated proteins were separated by SDS-PAGE and blotted for HMCES. Top panel, chromatin was directly analyzed. Bottom panel, recovered chromatin was treated with the deubiquitylating enzyme USP21 before SDS-PAGE. Non-specific bands are marked with asterisks. **d**, Left, experimental strategy to inhibit nascent strand approach. pICL^AP^ was replicated in NEIL3-depleted egg extract supplemented with p97 inhibitor for 60 min. Reactions were then treated with the polymerase inhibitor aphidicolin (or DMSO control) and rNEIL3 to allow for ICL unhooking. Right, chromatin was recovered under the indicated conditions and analyzed as in **c**.

Given the inhibitory effect of HMCES on TLS (Fig. 4), we expected that blocking SPRTN-dependent HMCES-DPC proteolysis would slow TLS. Surprisingly, while SPRTN-depletion inhibited TLS past a previously characterized HpaII model DPC^25^ (Extended Data Fig. 7a-d), it slightly accelerated TLS past the HMCES-DPC during AP-ICL repair (Extended Data Fig. 7a,b,e,f). This observation is consistent with a report that SPRTN can antagonize TLS by sequestering the TLS subunit POLD3^26^ and suggests that regulation of TLS by SPRTN depends on DPC identity and context. These results indicate that, although HMCES-DPC formation poses an impediment to AP site bypass, SPRTN-dependent proteolysis of HMCES is not a prerequisite for TLS past the cross-link.

**Fig. 7:**
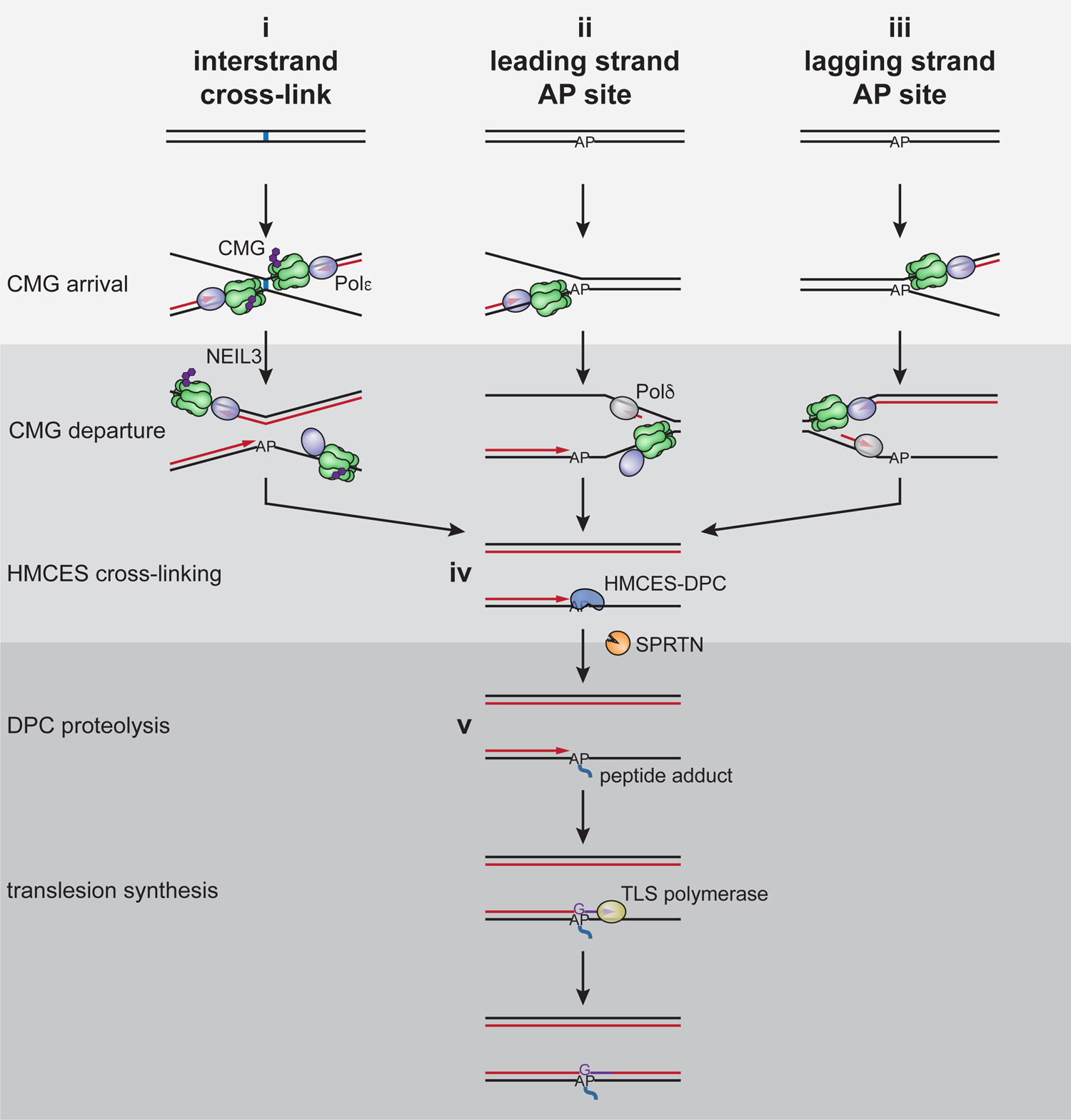
Model for AP site protection by HMCES during DNA replication. (i) Fork convergence at an ICL activates NEIL3-dependent unhooking and introduces an AP site in the leading strand template. The AP site is bypassed by CMG but stalls DNA polymerase, resulting in uncoupling of DNA unwinding from leading strand synthesis by Pol*ε*. This exposes the AP site at a ssDNA/dsDNA junction, where it is cross-linked by HMCES. (ii) AP sites encountered in the leading strand template due to spontaneous depurination/depyrimidination or incomplete BER are also bypassed by CMG. This again leads to CMG uncoupling and HMCES-DPC formation at a ssDNA/dsDNA junction. (iii) AP sites in the lagging strand template are bypassed without CMG uncoupling, and HMCES cross-links to the AP site embedded in ssDNA. Pol*δ* extends the lagging strand up to the HMCES-DPC, where upon synthesis stalls. (iv) In all cases, a ssDNA/dsDNA junction abutting the HMCES-DPC activates proteolysis by SPRTN. The HMCES-DPC and the peptide adduct generated after proteolysis stabilize the AP site until the lesion is bypassed by TLS (v) or an alternative, error-free mechanism (not depicted). HMCES-DPC formation introduces a bias for dG insertion opposite the AP site.

### Leading strand extension activates HMCES-DPC proteolysis

When the replisome collides with a HpaII DPC, CMG first bypasses the DPC, whereafter extension of the nascent leading strand up to the DPC triggers SPRTN-dependent proteolysis^23, 25^. Consistent with this observation, purified SPRTN preferentially targets DPCs positioned at a ssDNA-dsDNA junction^27^. We therefore tested whether degradation of the HMCES-DPC is also regulated by approach of the nascent DNA strand. pICL^AP^ was replicated in NEIL3-depleted extract supplemented with p97 inhibitor, which caused leading strand stalling 20 to 40 nucleotides from the ICL due to the footprint of the converged CMGs^15^ (Figs. 6d and 3b). rNEIL3 was then added back to activate ICL unhooking in the presence or absence of the replicative polymerase inhibitor aphidicolin, which blocked leading strand extension to the ICL^25, 28^. Chromatin was then recovered and blotted for HMCES. Compared to the DMSO control, aphidicolin greatly stabilized HMCES on chromatin (Fig. 6d, lanes 3-10). We conclude that SPRTN-dependent degradation of an HMCES-DPC is enhanced by approach of the leading strand, as seen during canonical DPC repair.

## Discussion

Current models envision that HMCES cross-links to pre-existing AP sites when these are encountered by the replication fork, but it has been unclear whether HMCES-DPC formation occurs as part of any DNA repair pathway. Here, we show that an HMCES-DPC is formed as a constitutive intermediate in the repair of DNA interstrand cross-links. We exploit this observation to elucidate the dynamics of HMCES cross-linking to DNA during replication and its effect on repair. Our data indicate that once the AP-ICL is unhooked by NEIL3 and CMG translocates past the newly formed AP site, HMCES cross-links to the AP site and protects it from cleavage. Subsequent extension of the leading strand to the HMCES-DPC triggers SPRTN-dependent HMCES proteolysis. Ultimately, the adduct is bypassed by TLS, introducing a bias for deoxyguanosine insertion opposite the AP site. Our results suggest a unified model to explain how HMCES is deployed during DNA replication to preserve genome stability.

### HMCES as a constitutive replication-coupled repair intermediate

Our observation that the HMCES-DPC is a bona fide intermediate in the NEIL3 ICL repair pathway raises the possibility that this structure participates in other DNA repair pathways. For example, the RPA2 winged helix domain was recently shown to stimulate excision of deoxyuracil from ssDNA by the UNG glycosylase in vitro, implying that RPA acts as a scaffold for a base excision repair mechanism that operates during replication^29^. It was further proposed that an HMCES-DPC could then stabilize the AP site produced by UNG and facilitate error-free bypass of the lesion. HMCES-DPC formation may therefore integrate multiple distinct replication-coupled repair pathways that generate AP site intermediates and enforce a common mechanism for bypassing various lesions.

### The timing and context of HMCES-DPC formation

Our data suggest that HMCES cross-links to the AP site after CMG has traveled beyond the unhooked ICL and before TLS occurs. Thus, we find that HMCES does not influence the rapid kinetics of leading strand extension to the AP site (Fig. 3), implying that CMG does not need to bypass an intact HMCES-DPC (Fig. 7, i and iv). Although we cannot rule out that CMG bypass of an HMCES-DPC is so fast that it does not delay leading strand extension, two additional lines of evidence suggest that HMCES-DPC formation is slower than CMG translocation past the AP site. First, the accumulation of HMCES-DPCs is delayed by several minutes relative to ICL unhooking (Fig. 1c, compare accumulation of unhooked strands with accumulation of adducted Top strand). Second, even in the presence of HMCES, a substantial fraction of recovered AP-ICL repair intermediates is initially susceptible to cleavage by exogenous APE1 (Extended Data Fig. 1e,f; compare 30- and 45-minute time points). Taken together, these data indicate that HMCES-DPC formation protects AP sites generated by ICL unhooking only after a delay of several minutes, whereas CMG departure after ICL unhooking should be immediate (Fig. 3). On the other hand, the HMCES-DPC clearly delays the already slow leading strand extension beyond the AP site (Fig. 4). Therefore, the HMCES-DPC probably forms after CMG departure but before TLS. Since leading strand extension up to the AP site should be almost instantaneous after ICL unhooking and CMG departure, we propose that HMCES cross-links the AP site at a ssDNA/dsDNA junction (Fig. 7, iv). This model is consistent with recent structural and biochemical studies using a bacterial SRAP domain, which suggest that an SRAP-DPC can accommodate a nascent strand at the AP site^4, 6, 7^. As seen for purified HMCES^3, 4^ (Extended Data Fig. 1b), HMCES in egg extracts can also cross-link to AP sites in a fully ssDNA region when nascent strand approach is blocked using aphidicolin (Fig. 3d).

HMCES cross-linking therefore exhibits a degree of flexibility that should allow it to protect AP sites encountered on either the leading or lagging strands.

### What causes AP site cleavage in the absence of HMCES?

A key question is what causes AP site cleavage and DSB formation in the absence of HMCES. AP site cleavage could occur either spontaneously (due to *β*-elimination) or via enzymatic action. Spontaneous *β*-elimination is unlikely to be the main source of AP site cleavage because the AP site remains intact during extensive workup of ICL repair intermediates (e.g., Extended Data Fig. 1c-f), suggesting that it is relatively stable. On the other hand, numerous enzymes present in egg extract can catalyze strand incision at an AP site. These enzymes include AP endonucleases such as APEX1, which cleaves the majority of AP sites in cells^30^. Although APEX1 has a strong preference for dsDNA AP sites, it has significant activity on ssDNA AP sites. Depletion of APEX1 from egg extracts did not reduce AP site cleavage during ICL repair (data not shown). Although cleavage could be mediated by APEX2, knocking down APEX1 and APEX2 in mammalian cells did not suppress DSB formation in the absence of HMCES^9^. ssDNA AP sites could also be cleaved by one or more AP lyases (DNA Pol*β*, PARP1, Ku, and the bifunctional DNA glycosylases NTH1, OGG1, NEIL1, NEIL2, and NEIL3). AP lyases catalyze strand incision at an AP site through Schiff base formation followed by *β*-elimination, a reaction that HMCES is expected to repress^4^. In the future, it will be important to identify the enzyme(s) that promote DSB formation in the absence of HMCES.

### Redundant mechanisms that suppress AP site cleavage?

While HMCES-DPC formation protected AP sites from cleavage during ICL repair, loss of HMCES did not lead to wide-spread DSB accumulation, even when AP site bypass was blocked (Figs. 2 and 3). This suggests that ssDNA AP sites are stabilized by one or more redundant mechanisms. ssDNA at converged replication forks is probably bound by RPA, which could non-specifically occlude AP endonuclease/lyase activity. A DNA polymerase, such as Pol *δ*, could also bind the 3’ end of nascent DNA at the AP site and occlude enzymatic AP site cleavage.

Alternatively, ssDNA AP sites might be protected by a more specialized DNA binding activity analogous to the Shu complex, which suppresses AP site-induced genome instability in yeast^31^. It will be interesting to determine whether an analogous Rad51 paralog complex contributes to AP site stability during DNA replication in vertebrates.

### The interplay of HMCES with lesion bypass

Previous reports indicate that HMCES antagonizes TLS and promotes error-free bypass of AP sites in human cells, although the underlying mechanism is unknown^3, 8, 9^. Our results show that while HMCES slows the kinetics of REV1-dependent AP site bypass, the adduct is eventually overcome by TLS, and there is no template switching. We speculate that the TLS impediment posed by HMCES helps promote error-free bypass in mammalian cells. However, error-free bypass does not occur in egg extracts, possibly because extracts lack a critical factor needed for this pathway or because small plasmids replicated in egg extract lack sufficient sister chromatid cohesion to enable template switching. Importantly, TLS past the AP-site position did not require SPRTN-dependent proteolysis of the HMCES-DPC (Extended Data Fig. 7), suggesting that TLS can accommodate an intact HMCES-DPC. This accommodation may be facilitated by structural features of the HMCES SRAP domain. The SRAP domain of YedK interacts with only seven nucleotides along ssDNA and the covalent thiazolidine linkage is formed by the extreme N-terminus of the protein^4, 6, 7^. HMCES may therefore be able to transiently undock from ssDNA while remaining covalently tethered to an AP site, effectively behaving as a peptide adduct that can be accommodated in a TLS polymerase active site.

Following TLS, the HMCES-DPC is no longer located at a ssDNA/dsDNA junction and therefore would not be processed by SPRTN. In this scenario, the HMCES-DPC may be removed by an alternative protease, such as the proteasome or DDI2 (see below).

Given that purines are more susceptible to spontaneous hydrolysis of the *N*-glycosyl bond (depurination) and oxidative damage requiring base excision repair, we anticipated that the HMCES-DPC might minimize mutagenesis by promoting pyrimidine insertion opposite an AP site. We were therefore surprised to find that HMCES-DPC formation introduces a bias for dG insertion opposite the adducted AP site. This HMCES-dependent bias for dG insertion during TLS may indicate an inherent thermodynamic constraint imposed on TLS polymerase during DPC bypass, akin to the “A rule” observed for insertion opposite an unprotected AP site. Alternatively, dG insertion bias may indicate that the SRAP domain evolved under conditions in which AP site formation was primarily driven by a process such as cytosine deamination. In this case a bias for dG insertion would help suppress the mutagenicity of AP sites.

### The mechanism of HMCES-DPC removal

We showed that during ICL repair in egg extract, HMCES-DPC proteolysis depended on SPRTN but not the proteasome. As seen for a model HpaII DPC, HMCES-DPC proteolysis by SPTRN depended on extension of the nascent leading strand up to the DPC, thereby placing the DPC at a ssDNA/dsDNA junction^25^ (Fig. 6d). Activation by nascent strand approach is therefore likely a general feature of SPRTN-dependent proteolysis, consistent with biochemical data indicating that SPRTN is specifically targeted to DPCs positioned at discontinuities in dsDNA^27^. In human cells, the proteasome inhibitor MG132 stabilizes HMCES on chromatin, suggesting that HMCES-DPCs can be degraded by the proteasome^3^. While the relative efficiencies of SPRTN- and proteasome-dependent proteolysis may differ in egg extracts and mammalian cells, proteasome inhibition may indirectly inhibit SPRTN-dependent HMCES proteolysis in cells. For example, proteasome inhibition might overwhelm the SPRTN-dependent pathway by increasing the total number of DPCs encountered during replication. Alternatively, because proteasome inhibition disrupts ubiquitin dynamics, it might also interfere with ubiquitin-dependent regulation of SPRTN activity^32^.

Interestingly, HMCES-DPCs accumulated as polyubiquitylated species that were still processed at a significant rate when both SPRTN and the proteasome were inhibited (Fig. 6c), suggesting the existence of an alternative HMCES-processing pathway. Recently, it was reported that *S. cerevisiae* Ddi1 (the homologue of vertebrate DDI2) removes a subset of DPCs in cells and that it can partially compensate for loss of the SPRTN homologue Wss1^33^.

Moreover, Ddi1 was also shown to be activated by long polyubiquitin chains *in vitro*^34^. Recent experiments performed in egg extracts have demonstrated that the E3 ubiquitin ligase RFWD3 polyubiquitylates an HMCES-DPC in an engineered ssDNA region of a circular plasmid ^35^.

Taken together, these observations suggest that RFWD3-dependent polyubiquitylation of an HMCES-DPC may induce DDI2-dependent proteolysis, although this possibility remains to be tested.

### A general model for HMCES action during DNA replication

Our data show that an HMCES-DPC is a constitutive intermediate of AP-ICL repair; it forms after CMG translocates beyond the unhooked ICL, and it protects the newly formed ssDNA AP site until after it is rendered double-stranded by TLS (Fig. 7, i). Based on this model, we infer that a similar mechanism operates when replication forks encounter AP sites generated by base hydrolysis or BER. We propose that when the AP site is located in the leading strand template, CMG passes over the lesion and leading strand synthesis stalls at the lesion (Fig. 7, ii). As seen during AP-ICL repair, HMCES cross-links to the AP site at a ssDNA/dsDNA junction. Although HMCES is immediately proteolyzed by SPRTN, the HMCES thiazolidine cross-link is sufficient to protect the AP site during the vulnerable period before TLS. A related mechanism would operate when forks encounter AP sites on the lagging strand template (Fig. 7 iii). As before, CMG rapidly translocates past the AP site, but due to the time required to initiate a new Okazaki fragment, the HMCES-DPC probably forms in a fully ssDNA context.

Nevertheless, the HMCES-DPC is likely to be short-lived because it will be degraded when extension of an Okazaki fragment triggers SPRTN-dependent HMCES degradation. In all cases, HMCES-DPC formation delays nascent strand extension past the AP site by TLS and potentially allows for engagement of an error-free bypass mechanism.

## Methods

All experiments were performed at least twice, with a representative result shown. Sequences of oligonucleotides mentioned in methods can be found in Supplementary Table 1. Next generation sequencing read counts described in Fig. 5 and Extended Data Fig. 5 can be found in Supplementary Table 2. No statistical methods were used to predetermine sample size.

Experiments were not randomized and the investigators were not blinded to allocation during experiments and outcome assessment. All unique materials are available from commercial suppliers or are available upon request from the authors.

### Preparation of oligonucleotide duplexes with site-specific interstrand cross-links

Site specific cross-links were prepared as previously described^13, 36^. To generate the Pt-ICL containing oligonucleotide duplex, 1 mM cisplatin was converted to activated monoaquamonochloro cisplatin by incubation in 10 mM NaClO_4_, 0.95 AgNO_3_ for 24 hours at 37 °C in the dark. Cisplatin monoadduct was then generated by incubating 0.125 mM Pt-ICL top oligonucleotide in 5.63 mM 10 mM NaClO_4_, 0.375 mM monoaquamonochloro cisplatin (in activation mixture) for 12 min at 37 °C. The reaction was quenched by addition of NaCl to 0.1 M. The monoadducted oligonucleotide was then purified with a Mono Q 5/50 GL column using a gradient from 370 mM to 470 mM NaCl in 10 mM NaOH over 40 column volumes.

Fractions containing the monoadducted oligonucleotide were pooled and adjusted to 2 mM MgCl_2_. 1.05 molar equivalents of Pt-ICL bottom oligonucleotide were added and buffer exchange was performed with 100 mM NaClO_4_ using an Amicon Ultra-15 3K filter unit at 4 °C. The oligonucleotides in 100 mM NaClO_4_ were allowed to cross-link by incubation at 37 °C for 48 hours. The cross-linked oligonucleotide duplex was then purified with a Mono Q 5/50 GL column using a gradient from 550 mM to 700 mM NaCl in 10 mM NaOH over 40 column volumes. Fractions containing the Pt-ICL duplex were pooled and buffer exchange was performed into 10 mM Tris-HCl (pH 7.4), 10 mM NaClO_4_ using an Amicon Ultra-15 3K filter unit at 4 °C. The cross-link was stored at −80 °C. To generate AP-ICL containing oligonucleotide duplexes, the appropriate complementary oligonucleotides (AP-ICL top and AP-ICL bottom; AP-ICL^rev^ top and AP-ICL^rev^ bottom; AP/A top and AP/A bottom; AP/G top and AP/G bottom; AP/T top and AP/T bottom; AP/C top and AP/C bottom) were annealed in 30 mM HEPES-KOH (pH 7.0), 100 mM NaCl by heating to 95°C for 5 min and cooling at 1°C/min to 18°C. The annealed duplex was then treated with uracil glycosylase (NEB) in 1x UDG buffer (20 mM Tris-HCl, 10 mM DTT, 10 mM EDTA [pH 8.0]) for 120 min at 37°C followed by extraction with phenol:chloroform:isoamyl alcohol (25:24:1; pH 8) and ethanol precipitation. The duplex was then dissolved in 50 mM HEPES-KOH (pH 7.0), 100 mM NaCl and incubated at 37°C for 5 to 7 days to allow cross-link formation. Cross-linked DNA was purified on a 20% polyacrylamide, 1x TBE, 8 M urea gel. The cross-linked products visualized by UV shadowing, eluted from crushed gel slices into TE (pH 8.0), extracted with phenol:chloroform:isoamyl alcohol (25:24:1; pH 8) and ethanol precipitated. The cross-links were dissolved in 10 mM Tris-HCl (pH 8.5) and stored at −80 °C.

### Preparation of plasmids containing cross-links (pICL)

Plasmids containing ICLs were prepared as described previously^13, 36, 37^. Briefly, the backbone plasmid (with or without 48 lacO repeats) was digested with BbsI in NEBuffer 2.1 for 24 hours at 37 °C followed by extraction with phenol:chloroform:isoamyl alcohol (25:24:1; pH 8) and ethanol precipitation. The linearized plasmid was dissolved in TE (pH 8.0) and purified over a HiLoad 16/60 Superdex 200 column using isocratic flow of TE (pH 8.0). Fractions containing the digested plasmid were pooled, ethanol precipitated and dissolved in 10 mM Tris-HCl (pH 8.5). The ICL-containing duplexes were ligated into the plasmid backbone using 400 U/mL NEB T4 DNA ligase in 1x ligase buffer (50 mM Tris-HCl (pH 7.5), 10 mM MgCl_2_, 1 mM ATP, 10 mM DTT) at room temperature for 24 hours. The ligation reactions were concentrated using Qiagen 500-tips and DNA was eluted with 1.25 M NaCl, 50 mM Tris-HCl (pH 8.5). The DNA was dialyzed into TE (pH 8.0) and concentrated with an Amicon Ultra-15 10K filter unit. CsCl was added to the DNA to a homogenous solution density of 1.6 g/mL and ethidium bromide was added to 50 µg/mL. The DNA was then transferred to a Quick-Seal tube and spun for 23 hours at 20 °C in an NVT-90 rotor at 75,000 rpm. The covalently closed circular plasmid was collected and extracted with an equal volume of saturated isobutanol to remove ethidium bromide. The DNA was then dialyzed into TE (pH 8.0), concentrated with an Amicon Ultra-15 10K filter unit, and snap frozen and stored at −80 °C. pICL^AP^ was used for the experiments shown in Fig. 1c-e, Fig. 2a,c, Extended Data Fig. 1c-f, Extended Data Fig. 2b,c, and Extended Data Fig. 3. pICL-lacO^AP^ was used for the experiments shown in Fig. 2d, Fig. 6b-d, Extended Data Fig. 2d-k, Extended Data Fig. 5c, and Extended Data Fig. 6. pICL-lacO^AP-rev^ was used for the experiments shown in Fig. 3b,c, Fig. 4a-c, Extended Data Fig. 4, and Extended Data Fig. 7b,e,f. pDPC^me^, used for the experiments shown in Extended Data Fig. 7b-d, was a gift from Justin Sparks and was prepared as described previously^23^.

### Preparation of Xenopus egg extracts

Animal work performed at Harvard Medical School was approved by the Harvard Medical Area Standing Committee on Animals (HMA IACUC Study ID IS00000051-3, approved 10/25/2017). The institution has an approved Animal Welfare Assurance (#A3431-01) from the NIH Office of Laboratory Animal Welfare. Animal work performed at Caltech was approved by the Institutional Animal Care and Use Committee (IACUC Protocol IA20-1797, approved 5/28/2020). The institution has an approved Animal Welfare Assurance (#D16-00266) from the NIH Office of Laboratory Animal Welfare. Preparation of high-speed supernatant (HSS) and nucleoplasmic extracts (NPE) from Xenopus laevis eggs was performed as described previously^38^. Briefly, HSS was prepared from eggs collected from six adult female frogs. Eggs were de-jellied in 1 L 2.2% cysteine, pH 7.7, washed with 2 L 0.5x Marc’s Modified Ringer’s solution (MMR; 2.5 mM HEPES-KOH [pH 7.8], 50 mM NaCl, 1 mM KCl, 0.25 mM MgSO_4_, 1.25 mM CaCl_2_, 0.05 mM EDTA), and washed with 1 L Egg Lysis Buffer (ELB; 10 mM HEPES-KOH [pH 7.7], 50 mM KCl, 2.5 mM MgCl_2_, 250 mM sucrose, 1 mM DTT, and 50 μg/mL cycloheximide). Eggs were then packed in 14 mL round-bottom Falcon tubes at 180xg using a Sorvall ST8 swinging bucket rotor. Eggs were supplemented with 5 μg/mL aprotinin, 5 μg/mL leupeptin, and 2.5 μg/mL cytochalasin B and then crushed by centrifugation at 20,000xg for 20 min at 4°C in a TH13-6×50 rotor using a Sorvall Lynx 4000 centrifuge.

The low-speed supernatant (LSS) was collected by removing the soluble extract layer and supplemented with 50 μg/mL cycloheximide, 1 mM DTT, 10 μg/mL aprotinin, 10 μg/mL leupeptin, and 5 μg/mL cytochalasin B. This extract was then spun in thin-walled ultracentrifuge tubes at 260,000xg for 90 min at 2°C in a TLS 55 rotor using a tabletop ultracentrifuge. Lipids were aspirated off the top layer, and HSS was harvested, aliquoted, snap frozen in liquid nitrogen, and stored at −80°C. NPE preparation also began with extracting LSS, except eggs were collected from 20 female frogs, and the volumes used to de-jelly and wash the eggs were doubled (2 L 2.2% cysteine, 4 L 0.5x MMR, and 2 L ELB). LSS was supplemented with 50 μg/mL cycloheximide, 1 mM DTT, 10 μg/mL aprotinin, 10 μg/mL leupeptin, 5 μg/mL cytochalasin B, and 3.3 μg/mL nocodazole. The LSS was then spun at 20,000xg for 15 min at 4°C in a TH13-6×50 rotor using a Sorvall Lynx 4000 centrifuge. The top, lipid layer was removed, and the cytoplasm was transferred to a 50 mL conical tube. ATP regenerating mix (2 mM ATP, 20 mM phosphocreatine, and 5 μg/mL phosphokinase) was added to the extract. Nuclear assembly reactions were initiated by adding demembranated Xenopus laevis sperm chromatin^39^ to a final concentration of 4,400/μL. After 75-90 min incubation, the nuclear assembly reactions were centrifuged for 3 min at 18,000xg at 4°C in a TH13-6×50 rotor using a Sorvall Lynx 4000 centrifuge. The top, nuclear layer was then harvested and spun at 260,000xg for 30 min at 2°C in a TLS 55 rotor using a tabletop ultracentrifuge. Finally, lipids were aspirated off the top layer, and NPE was harvested, aliquoted, snap frozen in liquid nitrogen, and stored at −80°C.

### Protein expression and purification

A PCR fragment containing the *Xenopus laevis* HMCES sequence with C-terminal FLAG tag was obtained by amplifying an IDT gBlock containing the HMCES coding sequence with primers HMCES F and HMCES R. NEBuilder HiFi DNA assembly master mix was used according to the manufacturer’s instructions to insert the HMCES PCR fragment into a pFastBac1 backbone that had been amplified with primers FB1-FLAG F and FB1-FLAG R. The sequence of pFastBac1-HMCES-FLAG was confirmed by Sanger sequencing. The C2A mutation was introduced by quick change mutagenesis using primers HMCES C2A F and HMCES C2A R and the sequence of pFastBac1-HMCES-FLAG C2A was confirmed by Sanger sequencing. The W321A L322A mutations were introduced by inverse PCR using primers HMCES W321A L322A F and HMCES W321A L322A R and the sequence of pFastBac1-HMCES-FLAG W321A L322A was confirmed by Sanger sequencing. Baculoviruses expressing the rHMCES-FLAG proteins were then prepared using the Bac-to-Bac system according to the manufacturer’s protocols.

HMCES-FLAG proteins were expressed in 250 mL suspension cultures of Sf9 cells (Expression Systems) by infection with baculovirus expressing rHMCES-FLAG for 72 hrs. Sf9 cells were collected and suspended in 8 mL lysis buffer (50 mM Tris-HCl [pH 7.5], 300 mM NaCl, 5% glycerol, 1x cOmplete protease inhibitors, 0.5 mM PMSF, 0.2% [v/v] Triton X-100). Cells were lysed by sonication and the soluble fraction was collected by spinning the lysate at 68,000xg for 1 hour. The soluble lysate was incubated with 200 µL anti-FLAG M2 affinity resign for 90 min at 4°C. The resin was washed once with 10 mL lysis buffer, twice with wash buffer (50 mM Tris-HCl [pH 7.5], 300 mM NaCl, 5% glycerol, 0.2% (v/v) Triton X-100), and three times with buffer 50 mM Tris-HCl [pH 7.5], 300 mM NaCl, 5% glycerol. HMCES-FLAG proteins were eluted from the resin with 50 mM Tris-HCl [pH 7.5], 300 mM NaCl, 5% glycerol, 100 µg/mL 3x FLAG peptide. Fractions containing rHMCES-FLAG proteins were dialyzed against 50 mM Tris-HCl (pH8.0), 150 mM NaCl, 10% glycerol, 10 mM DTT at 4°C overnight and then again for 4 hours at 4°C. Aliquots of protein were snap frozen and stored at −80°C.

Biotinylated LacI was expressed and purified essentially as described previously^40^. Briefly, LacI with a C-terminal AviTag and biotin ligase were co-expressed in T7 Express Cells supplemented with 50 µM biotin from pET11a[LacR-Avi] and pBirAcm (Avidity) vectors, respectively. AviTag-LacI and biotin ligase expression were induced with 1 mM IPTG in media supplemented with 50 μM biotin to ensure efficient biotinylation of the AviTag-LacI. Cell pellets were lysed for 30 minutes at room temperature in lysis buffer containing 50 mM Tris-HCl (pH 7.5), 5 mM EDTA, 100 mM NaCl, 1 mM DTT, 10% sucrose (w/v), 1x cOmplete protease inhibitors, 0.2 mg/mL lysozyme, 0.1% Brij 58. The insoluble fraction was pelleted by centrifugation at 21,300xg for 1 hour at 4 °C in an Eppendorf 5424R centrifuge. Chromatin-bound LacI was then suspended in 50 mM Tris-HCl (pH 7.5), 5 mM EDTA, 1 M NaCl, 30 mM IPTG, 1 mM DTT and released from the DNA by sonication followed by addition of polymin P to 0.03-0.06 % (w/v) at 4 °C. LacI was then precipitated with 37% ammonium sulfate, pelleted by centrifugation, and resuspended in buffer containing 50 mM Tris-HCl (pH 7.5), 1 mM EDTA, 2.6 M NaCl, 1 mM DTT, 1x cOmplete protease inhibitors. Next, biotinylated LacI was bound to with SoftLink avidin resin for 90 min at 4 °C, washed with 50 mM Tris-HCl (pH 7.5), 1 mM EDTA, 2.6 M NaCl, 1 mM DTT, 1x cOmplete protease inhibitors, and eluted with 50 mM Tris-HCl (pH 7.5), 100 mM NaCl, 1 mM EDTA, 5 mM biotin, 1 mM DTT. Pooled fractions containing LacI were buffer exchanged into 50 mM Tris-HCl (pH 7.5), 150 mM NaCl, 1 mM EDTA, 1 mM DTT using an Amicon Ultra-.5 3K filter unit. LacI aliquots were snap frozen in liquid nitrogen and stored at −80 °C.

*Xenopus laevis* rNEIL3-FLAG and rNEIL3-FLAG K60A were expressed and purified as described previously^13^. Briefly, baculoviruses expressing rNEIL3-FLAG were prepared using the Bac-to-Bac system (Thermo Fisher Scientific) according to the manufacturer’s protocols. rNEIL3-FLAG protein was expressed in 250 mL Sf9 insect cell cultures by infection with baculovirus expressing NEIL3–FLAG for 48 hours at 27 °C. Sf9 cells were collected and suspended in 10 mL lysis buffer (50 mM Tris-HCl [pH 7.5], 300 mM NaCl, 10% [v/v] glycerol, 1x cOmplete protease inhibitors, 0.5 mM PMSF, 0.2% [v/v] Triton X-100). Cells were lysed by sonication, and the soluble fraction was collected by spinning the lysate at 25,000 rpm in a Beckman SW41 rotor for 1 hour at 4 °C. The soluble lysate was incubated with 200 µL anti-Flag M2 affinity resin for 90 min at 4 °C. The resin was washed once with 10 mL Lysis Buffer, twice with 50 mM Tris-HCl (pH 7.5), 300 mM NaCl, 10% glycerol, 0.2% (v/v) Triton X-100), and three times with 50 mM Tris-HCl (pH 7.5), 300 mM NaCl, 10% glycerol. rNEIL3–FLAG protein was eluted from the resin with 50 mM Tris-HCl (pH 7.5), 300 mM NaCl, 10% glycerol, 100 μg/mL 3×FLAG peptide. Fractions containing rNEIL3–FLAG protein were pooled and dialyzed against 50 mM HEPES-KOH (pH 7.0), 300 mM NaCl, 1 mM DTT, 20% glycerol at 4 °C for 12 h and then dialyzed against 50 mM HEPES-KOH (pH 7.0), 150 mM NaCl, 1 mM DTT, 15% glycerol at 4 °C for 3 h. Aliquots of protein were snap frozen and stored at −80 °C. rSPRTN-FLAG and rSPRTN-FLAG E89Q proteins were a gift from Alan Gao and purified as described previously^25^. rUSP21 protein was a gift from Daniel Finley.

### Immunodepletions

Immunodepletions using antibodies against HMCES, SPRTN, REV1, and NEIL3 were performed as described^13, 25, 41^. Briefly, protein A Sepharose Fast Flow beads were washed in 1x PBS (137 mM NaCl, 2.7 mM KCl, 10 mM Na_2_HPO_4_, 1.8 mM KH_2_PO_4_) and then incubated with an appropriate volume of antibodies overnight at 4 °C (3 volumes for antibodies against HMCES, SPRTN, and NEIL3; 1 volume for antibodies against REV1). The beads were then washed twice with 1x PBS, once with ELB, twice with 500 mM NaCl in ELB, and thrice with ELB. Three rounds of immunodepletion were performed by adding 5 volumes of egg extract to 1 volume of beads and incubating on a rotating wheel at 4 °C for 60 min. In the case of REV1 depletion, two rounds were performed using the N-terminal antibody and one round was performed using the C-terminal antibody. Extracts were spun for 30 seconds at 2,500 rpm in a S-24-11-AT rotor using an Eppendorf 5430R centrifuge and the supernatants were collected without disturbing the beads.

### Replication reactions

Replication reactions were performed essentially as described previously^39^. Briefly, licensing was performed by incubating 7.5 ng/µL plasmid with HSS (supplemented with 3 µg/mL nocodazole, 20 mM phosphocreatine, 2 mM ATP, and 5 µg/mL creatine phosphokinase) for 30 min at room temperature (∼21°C). Where indicated, licensing mixes were supplemented with 111-333 nM 3000 Ci/mmol [*α*-^32^P]dATP. Replication was initiated by adding 1 volume licensing mix to 2 volumes NPE mix (50% NPE, 20 mM phosphocreatine, 2 mM ATP, and 5 µg/mL creatine phosphokinase, 4 mM DTT in ELB). Where indicated, replication reactions were supplemented with 100 nM rNEIL3-FLAG, 100-200 nM rHMCES-FLAG, 30 nM rSPRTN-FLAG, 200 µM NMS-873, 200 µM MG 262, 2 µM AZD6738, or 75 µM aphidicolin. For analysis on native agarose gels, reactions were stopped at indicated time points by mixing 1 μL of replication reaction mix with 6 μL replication stop buffer (8 mM EDTA, 0.13% phosphoric acid, 10% ficoll, 5% SDS, 0.2% bromophenol blue, 80 mM Tris-HCl, pH 8.0) and then digested with 2.5 mg/mL proteinase K for 60 min at 37 °C. The DNA was then resolved on 0.8% agarose, 1x TBE gels. The gels were dried and visualized by phosphorimaging on a Typhoon FLA 9500 phosphorimager (GE Healthcare). For analysis of nascent DNA strands and strand-specific Southern blots, the replication reactions were stopped at indicated time points by mixing 4 μL of replication reaction mix with 40 μL clear replication stop mix (50 mM Tris [pH 7.5], 0.5% SDS, 25 mM EDTA). Quenched reactions were then digested with 0.2 mg/mL RNaseA for 30 min at 37 °C and then with 2 mg/mL proteinase K for 60 min at 37 °C. Samples were adjusted to 200 µL with 10 mM Tris-HCl (pH 8.5), extracted twice with phenol:chloroform:isoamyl alcohol, extracted once with chloroform, and ethanol precipitated. The recovered DNA was suspended in 10 µL 10 mM Tris-HCl (pH 8.5) and stored at −20 °C. For replications analyzed by APE1 digestion, 2.4 µL recovered DNA was incubated for 3 hours at 37 °C with 4 U HincII and 4 U APE1, 1x NEBuffer 4 in a total of 4 µL. Digestions were stopped with 8 µL replication stop buffer and resolved on 0.8% agarose, 1x TBE gels.

### Nascent strand analysis

Nascent strand analysis was performed as described^18^. Briefly, plasmid DNA was recovered from replication reactions as described above and 3 µL DNA was incubated for 3 hours at 37 °C with 4 U AflIII, 8 U EcoRI, 4 U HincII, 4 U APE1, 4 U FspI, or 8 U AatII as indicated in a total volume of 5 µL. Digestions were performed in 1x NEBuffer 3.1 or 1x NEB CutSmart buffer. Digestion reactions were stopped by addition of 2.5 µL gel loading buffer 2 (95% Formamide, 18 mM EDTA and 0.025% SDS, 0.025% xylene cyanol, 0.025% bromophenol blue). Samples were heated at 75 °C for 3 min, snap cooled on ice and resolved on 7% acrylamide (37.5:1), 8 M urea, 0.8x GTG buffer (71 mM Tris, 23 mM taurine, 0.4 mM EDTA) sequencing gels. The gels were dried and visualized by phosphorimaging on a Typhoon FLA 9500 phosphorimager (GE Healthcare).

### Strand-specific Southern blotting

Strand-specific Southern blot was performed essentially as previously described^18^. Briefly, 3 µL plasmid DNA recovered from pICL^AP^ replication reactions (as described above) was incubated at 37 °C for 3 hours with 4 U AseI, 8 U XhoI, 1x NEBuffer 3.1 in a total volume of 5 µL. Digestion reactions were stopped by addition of 2.5 µL gel loading buffer 2. Samples were heated at 75 °C for 3 min, snap cooled on ice and resolved on 8% acrylamide (19:1), 8 M urea, 0.8x GTG buffer sequencing gels. The gels were transferred to filter paper and then the DNA was transferred to Hybond-XL membrane in 0.5x TBE at 0.4 A using a Transblot SD semi-dry transfer cell. The membrane was washed with 4x SSC for 5 min and then cross-linked with 120,000 µJ/cm^2^ 254 nm UV light. The membrane was incubated in 25 mL Ultrahyb buffer for at least 6 hours at 42 °C. The 85mer top strand probe and 87mer bottom strand probe were 5’ end radiolabeled by incubating 0.2 µM oligonucleotide with 0.2 µM 3,000 Ci/mmol [*γ*-^32^P]ATP and 2,000 U T4 DNA ligase in 1x NEB T4 DNA ligase buffer for 30 min at 37 °C in a total volume of 50 µL. The reaction was then heated to 65 °C for 20 min and passed through a Micro Bio-Spin Column with Bio-Gel P-6. The radiolabeled probe was added to the Ultrahyb buffer and incubated at 42 °C for 24 hours. The membrane was washed twice with 2x SSC, 0.1% SDS at 42 °C for 5 min. The membranes were visualized by phosphorimaging on a Typhoon FLA 9500 phosphorimager (GE Healthcare).

### Plasmid pull-down

Plasmid pulldowns to monitor the HMCES-DPC were performed essentially as previously described^25^. Briefly, streptavidin-coupled magnetic Dynabeads (10 µL per pull down) were washed twice with 50 mM Tris-HCl (pH 7.5), 150mM NaCl, 1mM EDTA (pH 8.0), 0.02% Tween-20. Biotinylated LacI (0.4 pmol per 1 µL beads) was added to the beads and incubated at room temperature for 40 min. The beads were then washed four times with DPC pull-down buffer (20 mM Tris pH 7.5, 150 mM NaCl, 2 mM EDTA pH 8, 0.5% IPEGAL-CA630) and then stored in the same buffer on ice until needed. At the indicated times 8 μL of replication reaction was quenched into 400 μL of DPC pull-down buffer on ice. After all of the timepoints were quenched, 10 μL of LacI-coated streptavidin Dynabeads were added to each sample and allowed to bind for 30 min at 4 °C on a rotating wheel. The beads were then washed three times with DPC pull-down buffer and then twice with Benzonase buffer (20 mM Tris pH 7.5, 20 mM NaCl, 2 mM MgCl2, 0.02% Tween-20). Beads were suspended in 7.5 μL Benzonase buffer containing 250-300 U Benzonase and 2.5 µM USP21. Beads were incubated for 1 hour at 37 °C on a rotating wheel. The supernatant was collected and mixed with 7.5 µL 2x Laemmli loading buffer and analyzed by immunoblotting.

### Immunoblotting

Samples in 1x Laemmli loading buffer (generally equivalent to 1 µL replication reaction or chromatin recovered from 8 µL plasmid pull-down) were resolved on 10% or 4-15% acrylamide Mini-PROTEAN or Criterion TGX precast gels (Bio-Rad) and transferred to PVDF membranes (Perkin Elmer). Membranes were blocked in 5% non-fat milk in 1x PBST for 1 hour at room temperature, rinsed several times with 1x PBST, and then incubated with antibody diluted in 1x PBST overnight at 4 °C with shaking. The membranes were washed with three times in 1x PBST for 10-20 minutes at room temperature. The membranes were then incubated with goat anti-rabbit horseradish peroxidase-conjugated secondary antibodies diluted in 5% non-fat milk in 1x PBST for 1 hour at room temperature. The membranes were washed three times in 1x PBST for 10-20 minutes at room temperature. The membranes were then incubated with SuperSignal West Dura or ProSignal Pico Spray chemiluminescence substrate for 1-4 minutes at room temperature and imaged using an Amersham Imager 600 (GE Healthcare) or ChemiDoc (Bio-Rad) imaging systems. Contrast was occasionally adjusted to improve visualization of bands.

### Denaturing gel shift assays

AP-ICL top oligonucleotide was 5’ end radiolabeled by incubating 0.2 µM oligonucleotide with 0.2 µM 3,000 Ci/mmol [*γ*-^32^P]ATP and 2,000 U T4 DNA ligase in 1x NEB T4 DNA ligase buffer for 30 min at 37 °C in a total volume of 50 µL. The reaction was then heated to 65 °C for 20 minutes and passed through a Micro Bio-Spin Column with Bio-Gel P-6. The labeled oligonucleotide was extracted with phenol:chloroform:isoamyl alcohol (25:24:1; pH 8) and precipitated in ethanol. The labeled oligonucleotide (∼5 pmol) was treated with uracil glycosylase (NEB) in 1x UDG buffer (20 mM Tris-HCl, 10 mM DTT, 10 mM EDTA [pH 8.0]) for 120 min at 37°C followed by extraction with phenol:chloroform:isoamyl alcohol (25:24:1; pH 8) and ethanol precipitation. 1 nM oligonucleotide was incubated with 50 nM rHMCES-FLAG protein in 10 mM Tris-HCl [pH 8.0], 50 mM NaCl, 10 mM MgCl_2_, 0.1 mg/mL BSA, 2 mM DTT for 1 hour at 37°C. 3 µL reaction was quenched into 15 µL 86% (v/v) formamide, 1x TBE, 20 mM EDTA, 0.25% (w/v) bromophenol blue and resolved on a 10% polyacrylamide (19:1), 8M Urea, 1x TBE gel. The gels were visualized by phosphorimaging on a Typhoon FLA 9500 phosphorimager (GE Healthcare).

### Preparation of sequencing libraries

Pooled pICL^AP-X^ plasmids were replicated in mock- or HMCES-depleted egg extract supplemented with rHMCES-FLAG for 3 hours at room temperature as described above. The 10 µL reactions were quenched with 100 µL clear replication stop mix (50 mM Tris [pH 7.5], 0.5% SDS, 25 mM EDTA). Quenched reactions were then digested with 0.2 mg/mL RNaseA for 30 min at 37 °C and then with 2 mg/mL proteinase K for 60 min at 37 °C. Samples were adjusted to 400 µL with 10 mM Tris-HCl (pH 8.5), extracted twice with 400 µL phenol:chloroform:isoamyl alcohol, extracted once with 400 µL chloroform, and ethanol precipitated. The recovered DNA was suspended in 22 µL 10 mM Tris-HCl (pH 8.5). The DNA was then digested with 5 U Nt.BstNBI in 25 µL 1x NEBuffer 3.1 for 1 hour at 55 °C and then for 20 minutes at 80 °C. Samples were adjusted to 150 µL with 10 mM Tris-HCl (pH 8.5), extracted twice with 150 µL phenol:chloroform:isoamyl alcohol, extracted once with 150 µL chloroform, and ethanol precipitated. The recovered DNA was suspended in 8 µL 10 mM Tris-HCl (pH 8.5). DNA (∼10 ng) was amplified in 100 µL containing 1x NEB Phusion buffer, 0.2 mM each dNTP, 2.5 µL RA302 primer, 2.5 µM RA303 primer, and 2 U NEB Phusion polymerase. Amplification reactions were incubated at 98 °C for 30 seconds followed by 18 cycles of incubation at 98 °C for 10 seconds, 55 °C for 30 seconds, and 72 °C for 30 seconds, and then incubated at 72 °C for 10 minutes. The amplified products were inspected on a 1% agarose, 1x TBE gel stained with Sybr Gold, purified using the QIAquick PCR purification kit according to the manufacturer’s instructions, and submitted for next generation amplicon sequencing by Genewiz.

### Data availability statement

All data supporting the findings of this study are available within the article and its supplementary information files. Unprocessed and uncropped gel and blot images are available from the corresponding authors upon request.

---

**Extended Data Fig. 1:**
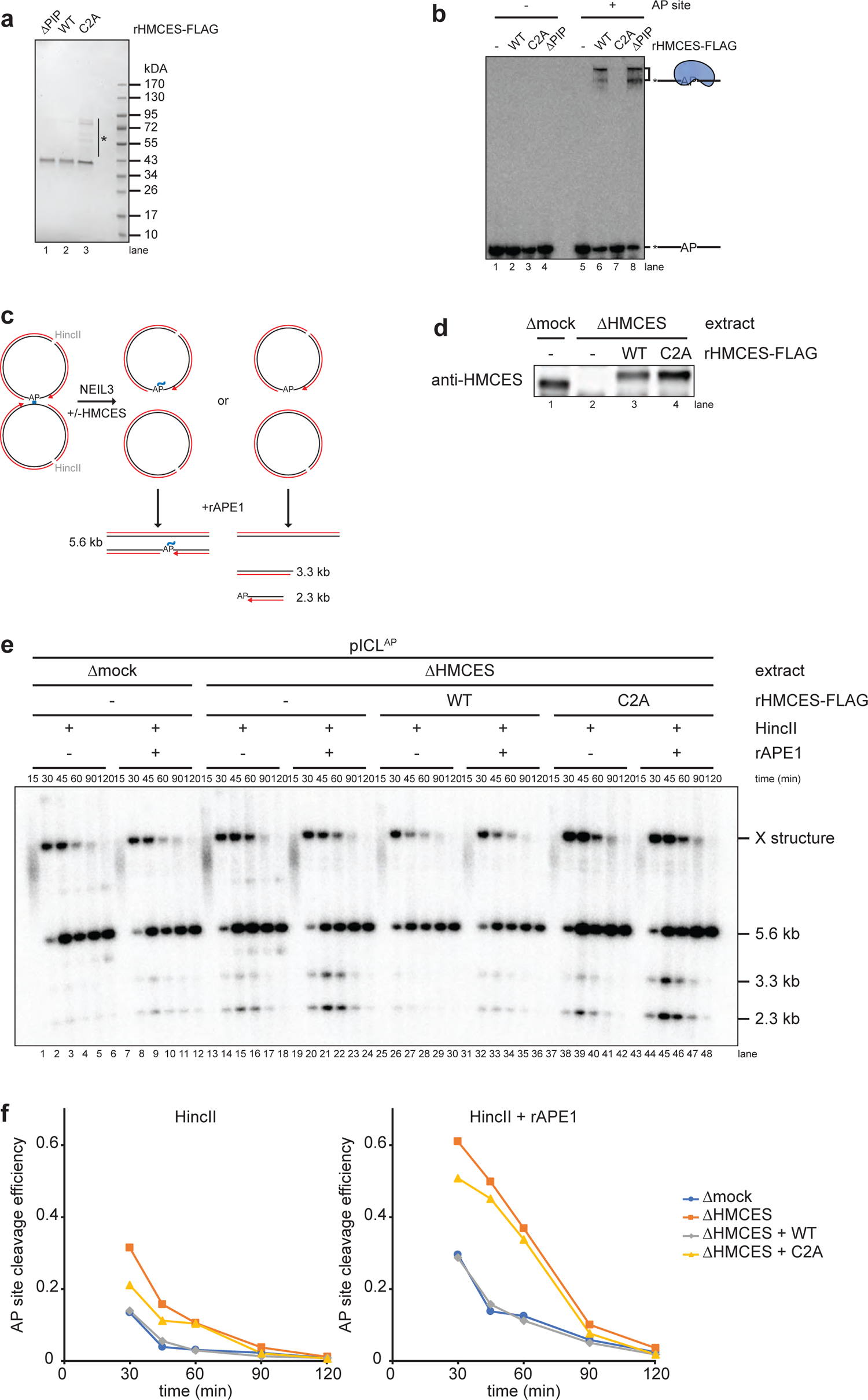
An HMCES-DPC shields AP sites. **a**, Purified recombinant FLAG-tagged *Xenopus laevis* HMCES proteins were resolved by SDS-PAGE and visualized by staining with InstantBlue. rHMCES^ΔPIP^ harbors W321A and L322A mutations that disrupt a conserved PIP-box that was previously show to mediate interaction with PCNA. Asterisk, contaminating bands. **b**, A 5’ end radiolabeled 20mer oligonucleotide with a single deoxyuracil (-AP site) or AP site (+AP site) was incubated with rHMCES proteins shown in (a) for 60 min. Samples were then resolved on a denaturing polyacrylamide gel and visualized by autoradiography. Reactions contained 1 nM oligonucleotide and 50 nM rHMCES. **c**, Schematic of species produced by digestion of pICL^AP^ replication intermediates with HincII and APE1. Digestion with HincII generates a 5.6 kb linear plasmid species while additional AP site cleavage by APE1 is expected to generate 2.3 kb and 3.3 kb species. **d**, HMCES immunodepletion. The extracts used in the reactions shown in **e** were blotted for HMCES. **e**, pICL^AP^ was replicated with [*α*-32P]dATP in the indicated egg extracts (shown in **d**). Samples were treated with proteinase K, phenol:chloroform extracted, and digested with HincII or with HincII and APE1. Digested DNAs were resolved on a native agarose gel and visualized by autoradiography. X structures indicate HincII-digested plasmids before ICL unhooking. **f**, Quantification of APE1 cleavage efficiency for the reactions shown in **e**. Cleavage efficiency was quantified as the intensity (Int) of 2.3 kb and 3.3 kb fragment bands in each lane divided by the total intensity of linear species bands ([Int^2.3kB^ + Int^3.3kB^]/[Int^2.3kB^ + Int^3.3kB^ + Int^5.6kB^]). The efficiency of HMCES-DPC formation in mock-depleted extract was estimated by subtracting the extent of rAPE1 cleavage in mock-depleted extract from the extent of cleavage in HMCES-depleted extract at 45 min (to maximize the absolute signal resulting from rAPE1 cleavage).

**Extended Data Fig. 2:**
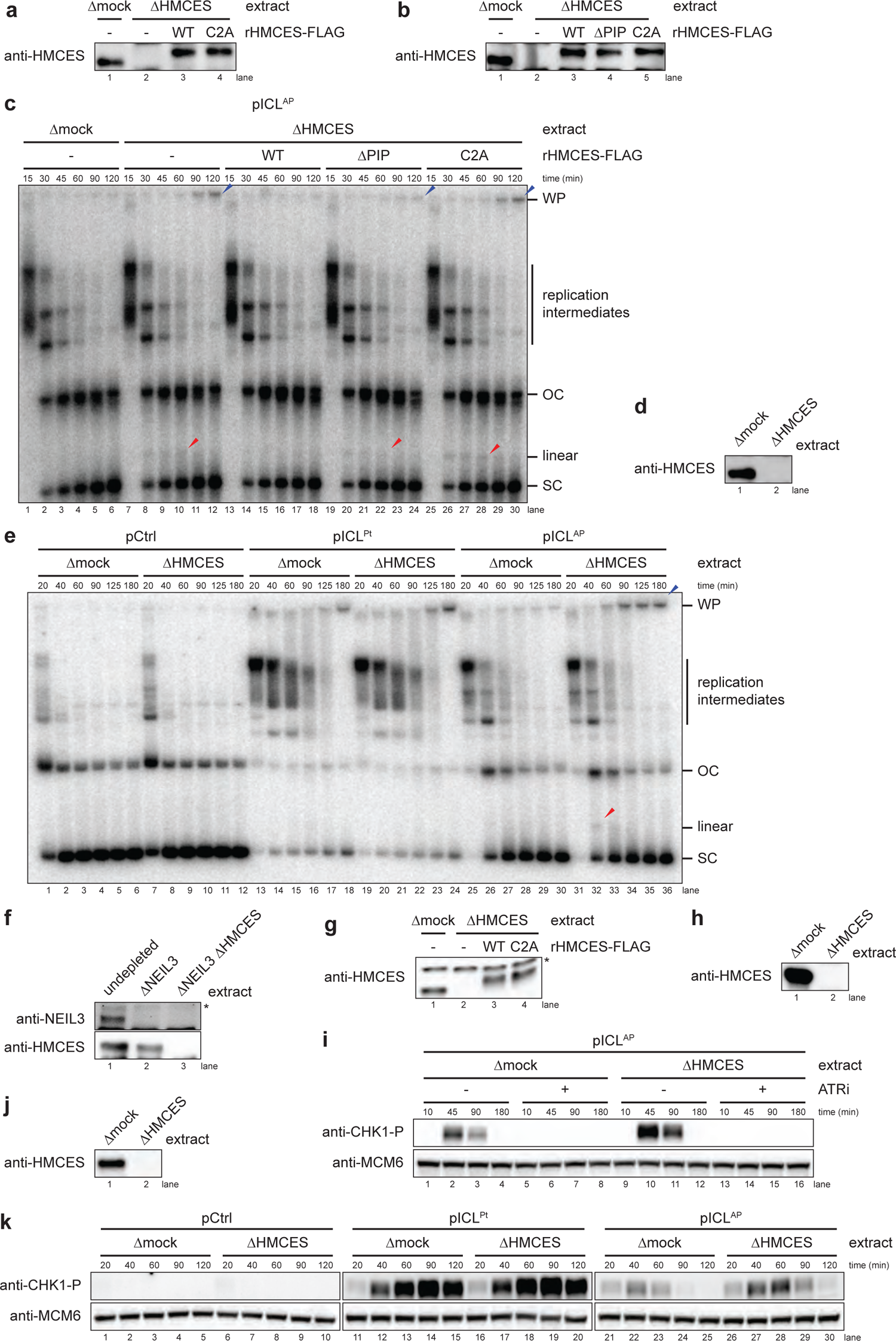
HMCES suppresses DSB formation specifically during NEIL3-dependent ICL repair. **a**, **b**, **d**, **f**, **g**, **h**, and **j**, HMCES and NEIL3 immunodepletions. The extracts used in Fig. 2a (**a**), Extended Data Fig. 2c (**b**), Extended Data Fig. 2e (**d**), Fig. 2c (**f**), Fig. 2d, (**g**), Extended Data Fig. 2i (**h**), and Extended Data Fig. 2k (j) were blotted for HMCES and NEIL3 as indicated. **c**, pICL^AP^ was replicated with [*α*-^32^P]dATP in mock- or HMCES-depleted extracts supplemented with rHMCES, as indicated. Replication intermediates were analyzed as in Fig. 2a. **e**, pCtrl, pICL^Pt^, or pICL^AP^ were replicated with [*α*-^32^P]dATP in mock- or HMCES-depleted extracts, as indicated. Replication intermediates were analyzed as in Fig. 2a. **i**, pICL^AP^ was replicated in mock- or HMCES-depleted extracts supplemented with ATR inhibitor AZD6738, as indicated. Replication reactions were separated by SDS-PAGE and blotted for phospho-CHK1 and MCM6 (loading control). Accumulation of phosphorylated CHK1 was blocked by AZD6738 treatment, indicating that CHK1 phosphorylation is dependent on ATR. **k**, pCtrl, pICL^Pt^, or pICL^AP^ was replicated in mock- or HMCES-depleted extracts, as indicated. Replication reactions were analyzed as in **i**. HMCES depletion increases the accumulation of phosphorylated CHK1 only during replication of pICL^AP^, indicating a specific role for HMCES in NEIL3-dependent ICL repair.

**Extended Data Fig. 3:**
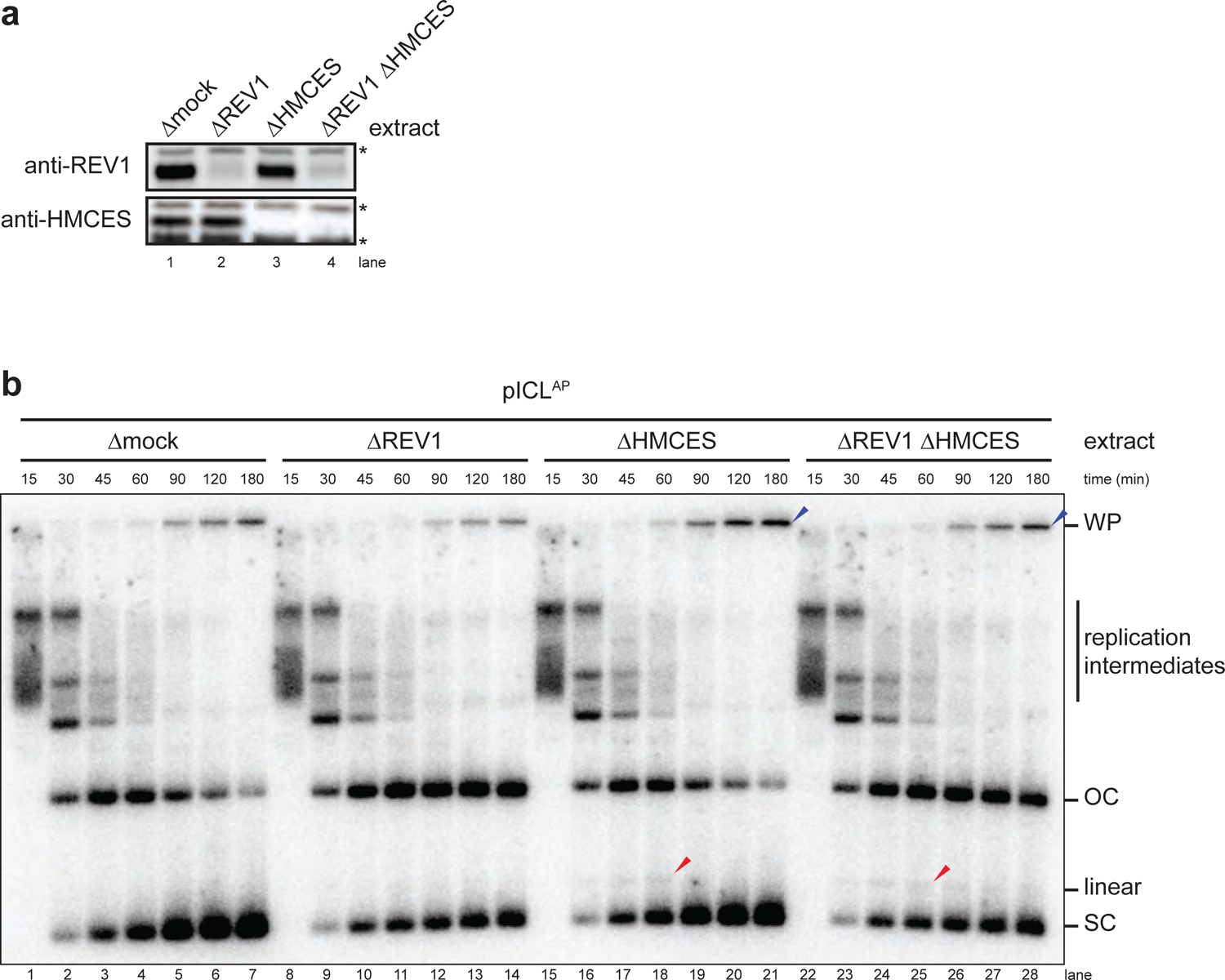
TLS inhibition does not enhance DSB formation during NEIL3-dependent ICL repair. **a**, REV1 and HMCES immunodepletion. The extracts used in the replication reactions shown in **b** were blotted for REV1 and HMCES. Asterisks, non-specific bands. **b**, pICL^AP^ was replicated with [*α*-^32^P]dATP in the indicated extracts (shown in **a**). Replication intermediates were analyzed as in Fig. 2a.

**Extended Data Fig. 4:**
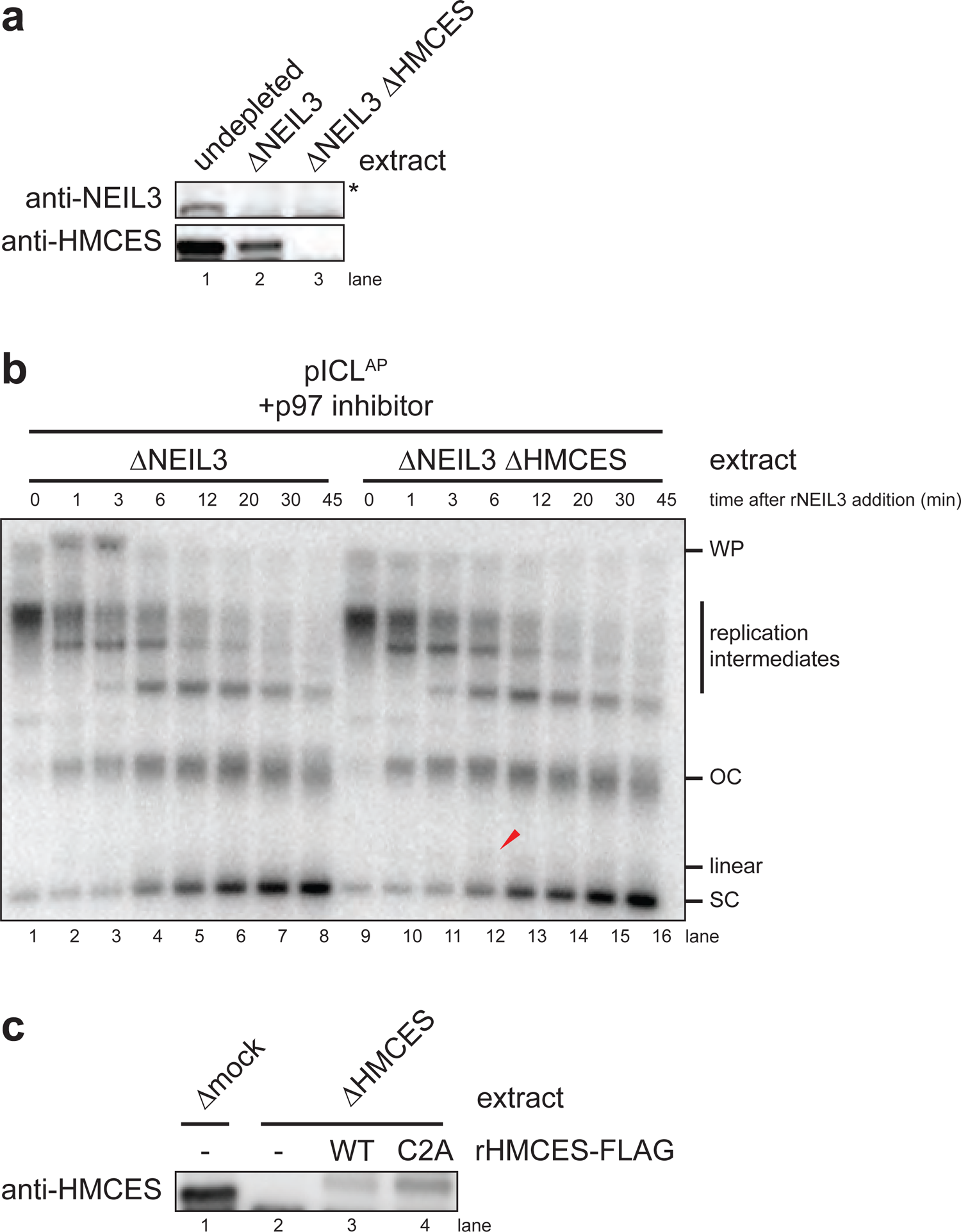
Synchronization of ICL unhooking by NEIL3-depletion and add back. **a**, NEIL3 and HMCES immunodepletion. The extracts used in the replication reactions shown in Fig. 3b were blotted for HMCES and NEIL3. Asterisk, non-specific band. **b**, pICL^AP^ was replicated in the presence of [*α*-^32^P]dATP in the indicated extracts (shown in **a**) supplemented with p97 inhibitor for 60 min. rNEIL3 was then added to the reactions to allow ICL unhooking. Replication intermediates were analyzed as in Fig. 2a. **c**, HMCES immunodepletion. The extracts used in Fig. 4 were blotted for HMCES.

**Extended Data Fig. 5:**
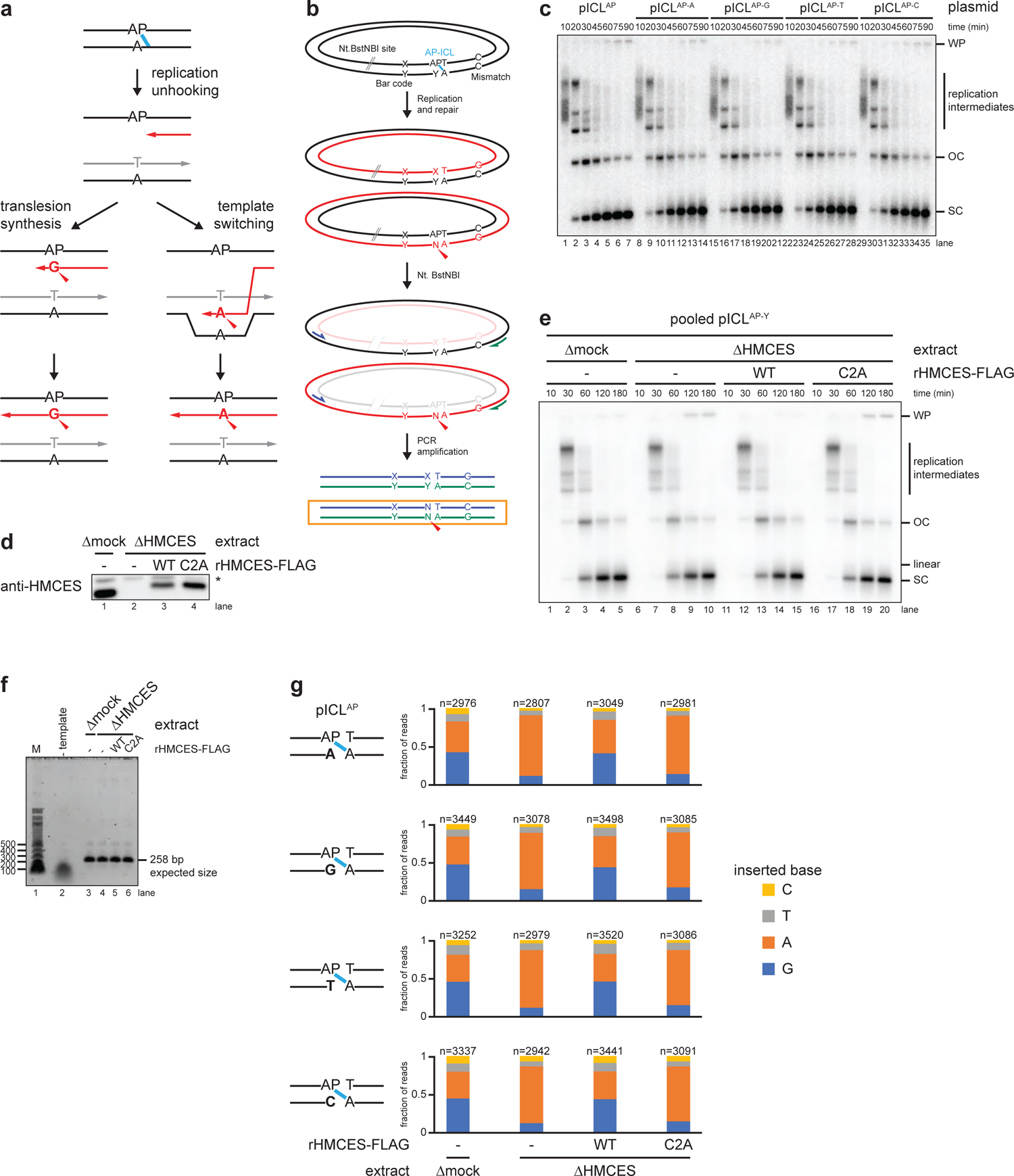
Nucleotide identity opposite the AP site has no effect on NEIL3-dependent ICL unhooking or TLS. **a**, Model of two alternative mechanisms of AP site bypass. Translesion synthesis (left branch) uses a specialized TLS polymerase for untemplated insertion of a nucleotide opposite the non-coding AP site. The nucleotide opposite the AP site in the parental plasmid does not influence insertion by the TLS polymerase. Template switching (right branch) uses the newly synthesized DNA of the other sister chromatid as a template for error-free bypass of the AP site. The inserted nucleotide is expected to co-vary with the nucleotide opposite the AP site in the parental plasmid. **b**, Plasmid design for next generation sequencing. We prepared four different plasmids, each containing an AP-ICL and a different nucleotide positioned opposite the AP site (Y), as well as a unique barcode. An Nt.BstNBI restriction site allows specific cleavage of the AP site-containing strand to prevent its amplification during PCR. A dC-dC mismatch allows reads produced from amplification of the nascent strand that has bypassed the AP site (PCR product in orange box) to be distinguished from reads derived from the corresponding parental strand (upper PCR product). **c**, The four AP-ICL plasmids, each containing a different nucleotide opposite the AP-site (described in **b**), were replicated in egg extract supplemented with [*α*-^32^P]dATP. Replication intermediates were analyzed as in Fig. 2a. The base opposite the AP site had no apparent effect on replication or repair efficiency. **d**, HMCES immunodepletion. The extracts used to generate the sequencing libraries described in Fig. 5 were blotted for HMCES. Asterisk, non-specific band. **e**, The four AP-ICL plasmids described in **b** were pooled and replicated with [*α*-^32^P]dATP in the same extracts (shown in panel d) used to generate sequencing libraries described in Fig. 5. Replication intermediates were analyzed as in Fig. 2a. **f**, Analysis of PCR amplicons used for sequencing. In parallel to the reactions shown in **e**, pooled pICL^AP^ plasmids were replicated in the indicated extracts (shown in **d**) supplemented with rHMCES, as indicated. DNA was extracted and digested with Nt.BstNBI (to cleave AP site-containing strands). The region of the replicated plasmids surrounding the ICL was then amplified by PCR. PCR amplicons were resolved by native agarose gel electrophoresis and visualized by Sybr Gold staining. **g**, Analysis of sequencing reads derived from individual pICL^AP^ plasmids. The barcode position was used to distinguish sequencing reads derived from the four different AP-ICL containing plasmids described in **b**. For each extract condition, we obtained >30,000 mapped reads, the vast majority (87.4%-87.8%) of which either perfectly matched the reference sequence or had a single point mutation corresponding to the position opposite the AP site. Of these reads, >11,000 in each condition derived from the nascent DNA stand produced upon bypass of the AP site. The fraction of reads corresponding to insertion of a given nucleotide opposite the AP site are plotted for each plasmid and extract condition. n, number of pooled nascent strand reads obtained for each condition. The result shows that the nucleotide opposite the AP site in the parental plasmid template does not influence the distribution of nascent DNA nucleotides inserted opposite the AP site after unhooking. Next generation sequencing read counts can be found in Supplementary Table 2.

**Extended Data Fig. 6:**
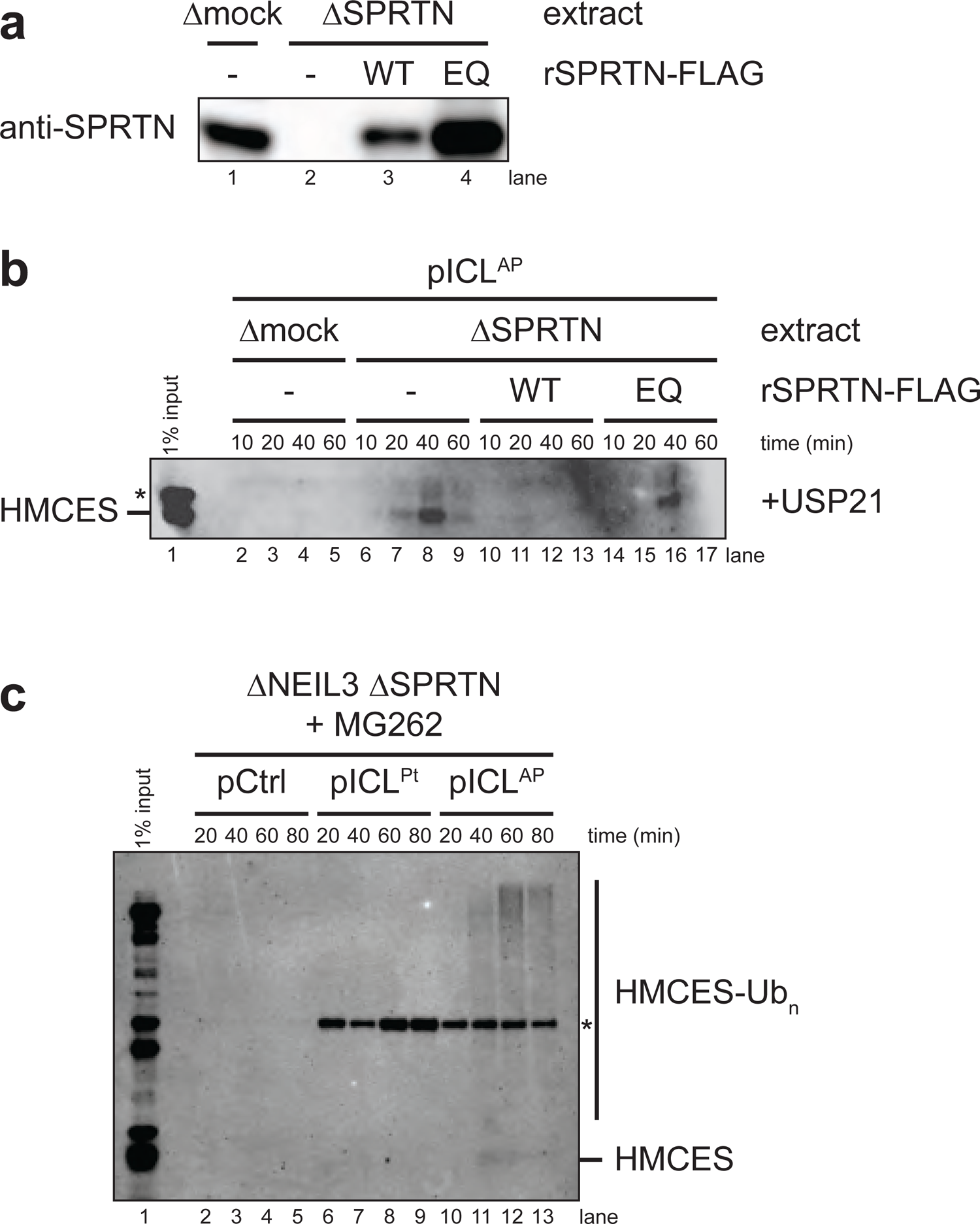
SPRTN protease activity is required for HMCES removal. **a**, SPRTN immunodepletion. The extracts used in the replication reactions shown in **b** were blotted for SPRTN. **b**, pICL^AP^ was replicated in mock- or SPRTN-depleted egg extract supplemented with wild-type (WT) or catalytically defective E89Q-mutated (EQ) rSPRTN, as indicated. Chromatin was recovered under stringent conditions, treated with the deubiquitylating enzyme USP21, and associated proteins were separated by SDS-PAGE and blotted for HMCES. **c**, pCtrl, pICL^Pt^, or pICL^AP^ were replicated in SPRTN-depleted egg extract supplemented with proteasome inhibitor MG262. Chromatin was analyzed as in Fig. 6c. Chromatin-associated HMCES is only observed in the pICL^AP^ replication reaction, implying a specific role for HMCES in NEIL3-dependent ICL repair.

**Extended Data Fig. 7:**
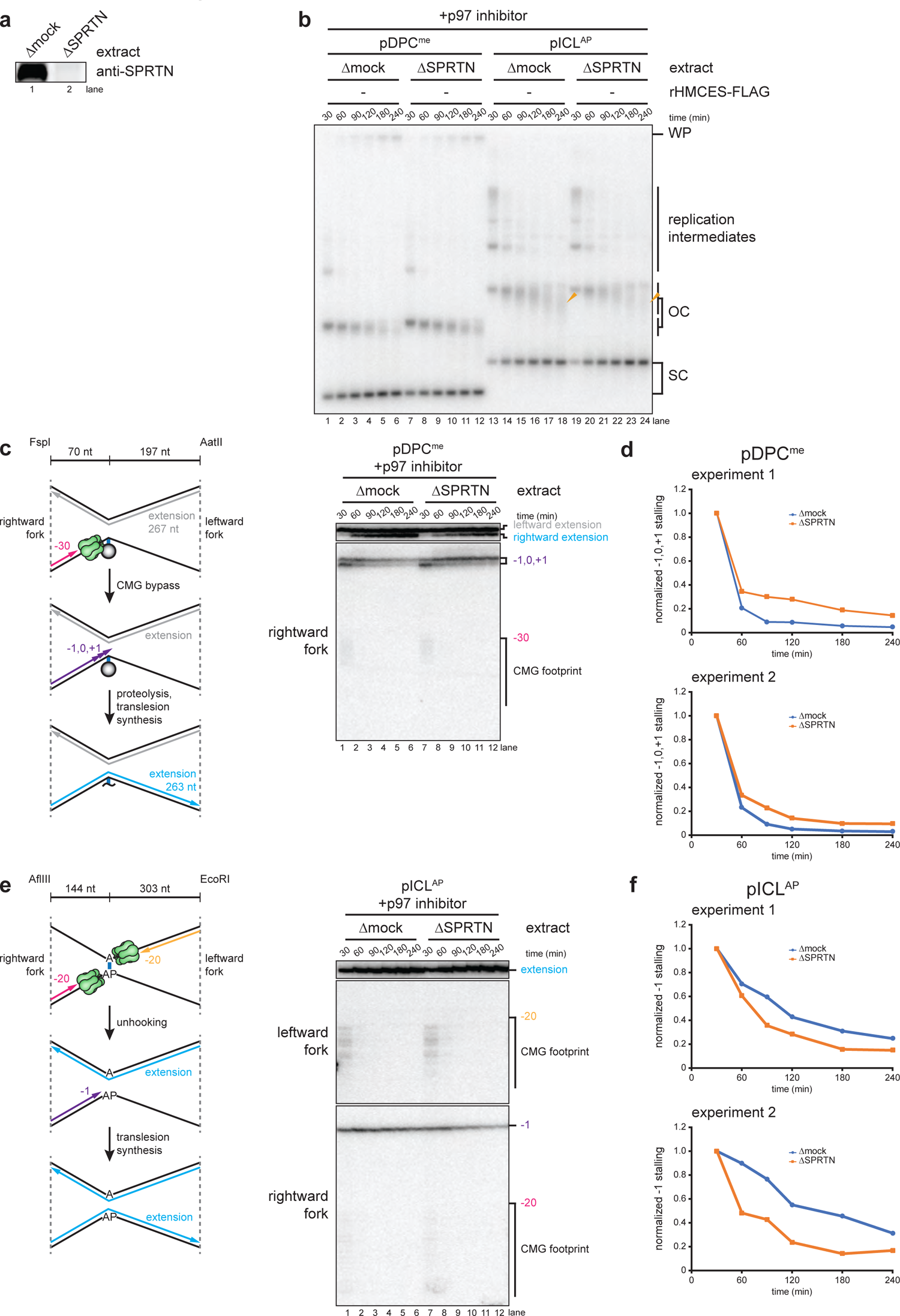
SPRTN-dependent proteolysis of the HMCES-DPC is not required for TLS past the AP site. **a**, SPRTN immunodepletion. The extracts used in **b**-**f** were blotted for SPRTN. **b**, A plasmid containing a methylated HpaII DPC that is refractory to degradation by the proteasome (pDPC^me^) or pICL^AP^ were replicated with [*α*-^32^P]dATP in mock- or SPRTN-depleted egg extracts supplemented with p97 inhibitor. Replication intermediates were analyzed as in Fig. 4a. **c**, Left, schematic of nascent strands generated during DPC repair. FspI and AatII cut 70 nucleotides to the left and 197 nucleotides to the right of the ICL, respectively, generating characteristic −30 stall, −1 to +1 stall, and strand extension products. Right, pDPC^me^ was replicated as in **b** and nascent DNA strands were isolated, digested with FspI and AatII, and resolved by denaturing polyacrylamide gel electrophoresis. As previously reported, SPRTN-depletion delays TLS past the methylated HpaII DPC, as evidenced by the persistence of rightward fork −1, 0, and +1 stall products and a delay in formation of rightward fork extension products. **d**, The persistence of the rightward fork −1, 0, and +1 stall products in **c** was quantified by dividing the summed intensity of the −1, 0, and +1 stall product bands in each lane by the intensity of the full-length rightward extension product band. Quantifications were normalized to the accumulation of the −1, 0, and +1 stall products at the 0 min time timepoint. Quantifications from two independent experiments are shown. **e**, Left, schematic of nascent strands generated during AP-ICL repair, as in Fig. 4b. Right, pICL^AP^ was replicated as in **b** and nascent DNA strands were analyzed as in Fig. 4b. In contrast to TLS past the methylated HpaII-DPC shown in **c**, SPRTN-depletion accelerated TLS past the HMCES-DPC, as evidenced by the faster disappearance of rightward leading strand −1 stall products. HMCES-depletion further accelerates disappearance of the rightward leading strand − 1 stall products. This result indicates that TLS past the AP site does not require SPRTN-dependent proteolysis of the HMCES-DPC formed during NEIL3-dependent ICL repair. **f**, The persistence of the rightward fork −1 stall product in **e** was quantified by dividing the intensity of the −1 stall product band in each lane by the intensity of the full length extension product band. Quantifications were normalized to the accumulation of the −1 stall product at the 0 min time timepoint. Quantifications from two independent experiments are shown.

**Supplementary Table 1.**
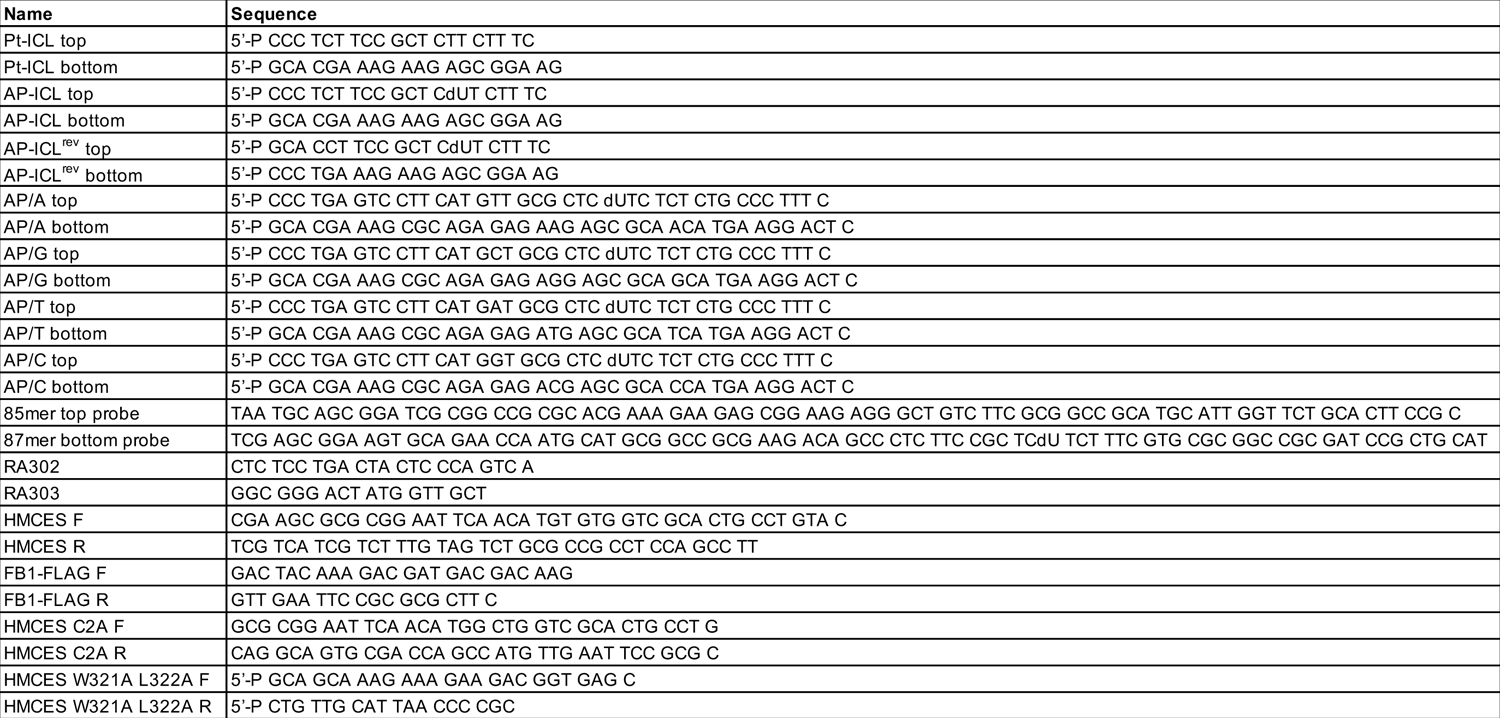
Oligonucleotide Sequences

